# Compound stimuli reveal the structure of visual motion selectivity in macaque MT neurons

**DOI:** 10.1101/692533

**Authors:** Andrew D Zaharia, Robbe L T Goris, J Anthony Movshon, Eero P Simoncelli

**Affiliations:** Center for Neural Science, New York University, New York, NY 10003; Howard Hughes Medical Institute, New York, NY 10003

## Abstract

Motion selectivity in primary visual cortex (V1) is approximately separable in orientation, spatial frequency, and temporal frequency (“frequency-separable”). Models for area MT neurons posit that their selectivity arises by combining direction-selective V1 afferents whose tuning is organized around a tilted plane in the frequency domain, specifying a particular direction and speed (“velocity-separable”). This construction explains “pattern direction selective” MT neurons, which are velocity-selective but relatively invariant to spatial structure, including spatial frequency, texture and shape. Surprisingly, when tested with single drifting gratings, most MT neurons’ responses are fit equally well by models with either form of separability. However, responses to plaids (sums of two moving gratings) tend to be better described as velocity-separable, especially for pattern neurons. We conclude that direction selectivity in MT is primarily computed by summing V1 afferents, but pattern-invariant velocity tuning for complex stimuli may arise from local, recurrent interactions.

**Significance Statement:** How do sensory systems build representations of complex features from simpler ones? Visual motion representation in cortex is a well-studied example: the direction and speed of moving objects, regardless of shape or texture, is computed from the local motion of oriented edges. Here we quantify tuning properties based on single-unit recordings in primate area MT, then fit a novel, generalized model of motion computation. The model reveals two core properties of MT neurons — speed tuning and invariance to local edge orientation — result from a single organizing principle: each MT neuron combines afferents that represent edge motions consistent with a common velocity, much as V1 simple cells combine thalamic inputs consistent with a common orientation.

## Introduction

Most neurons in extrastriate area MT (V5) are tuned for the speed and direction of visual motion (Dubner and Zeki, 1971; Van Essen et al., 1981; Maunsell and Van Essen, 1983), and many of them are selective for the coherent motion of complex patterns (Movshon et al., 1985). Such tuning is absent from the earliest stages of visual processing in primates, the retina and lateral geniculate nucleus. There, incoming visual signals are filtered without regard to direction, and are approximately separable in space and time (Enroth-Cugell et al., 1983; Derrington and Lennie, 1984). Motion-selective simple cells in primary visual cortex (V1) are tuned for motion in a manner that treats spatial and temporal frequency roughly separably (Tolhurst and Movshon, 1975), while a quarter of V1 complex cells treat them jointly (Priebe et al., 2006), consistent with speed tuning. V1 neurons provide input to MT, where neurons also tend to be speed tuned (Perrone and Thiele, 2001; Priebe et al., 2003).

Motion-selective V1 neurons are also orientation-selective, and their responses confound the direction of motion and the orientation of moving stimuli. In particular, they respond independently to each oriented component rather than to the pattern as a whole (Movshon et al., 1985). Under many conditions, humans perceive such complex patterns as moving coherently in a single direction (Wallach, 1935; Adelson and Movshon, 1982). Similarly, MT neurons signal coherent pattern motion, with some neurons being completely invariant to component orientation (Movshon et al., 1985). The degree to which MT neurons respond to the motion of individual components or the whole pattern lies on a continuum, quantified by a “pattern index” (see figure 1(a-c), Methods, and Movshon et al. (1985)).

**Figure 1.**
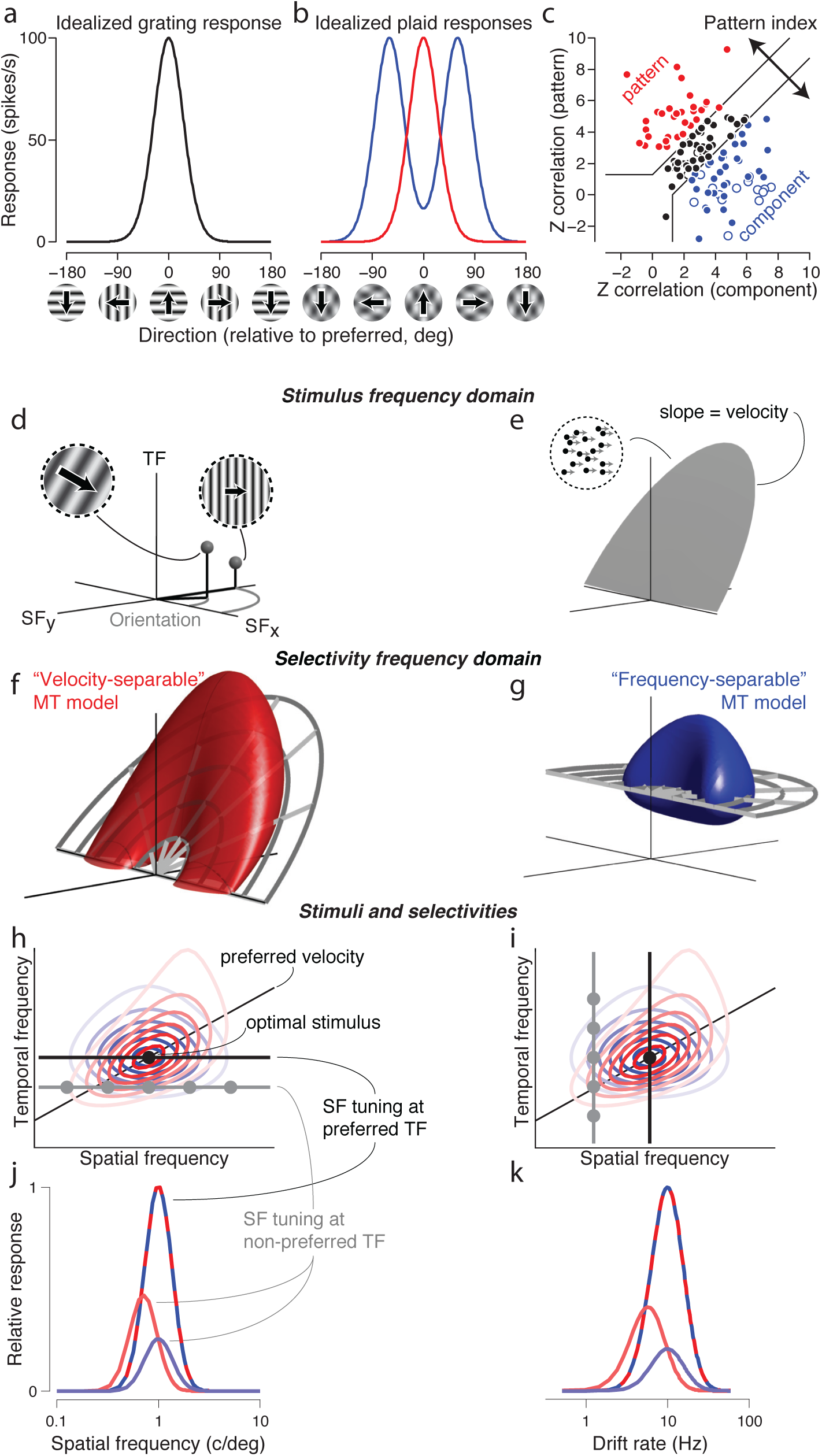
Pattern index, frequency-separable and velocity-separable hypotheses, and their predicted tuning. (a) Tuning curve of an idealized direction-selective neuron, responding to drifting gratings. (b) An ideal pattern-selective neuron exhibits a unimodal tuning curve for drifting plaids (red), while an ideal component neuron shows a bimodal tuning curve (blue). The peaks of the component neuron tuning curve correspond to the directions of the two gratings that comprise the plaid. (c) The “pattern index” captures the degree to which a given neuron is pattern or component selective. Each point represents the correlation of a given neuron’s measured tuning curve with the ideal component and pattern tuning (abscissa and ordinate, respectively; see methods for details), as predicted from its actual grating responses. Open and filled points correspond to neurons featured in this paper in V1 (*n* = 21) and MT (*n* = 112), respectively. (d) Three-dimensional frequency domain representation of moving images, with two spatial frequency axes and one temporal frequency axis. These coordinates can alternatively be expressed as orientation, spatial frequency, and temporal frequency. A single point in the frequency domain represents a single drifting sinusoidal grating. (e) The motion of a rigidly translating pattern (e.g., a field of dots moving with the same velocity) contains frequency components that lie on a plane through the origin. (f-g) Two possible hypotheses for MT selectivity in the frequency domain. In a velocity-separable receptive field (f), spatial and temporal frequency tuning are concentrated along a tilted, preferred velocity plane. In the frequency-separable prediction (g), spatial and temporal frequency tuning are independent. Note the velocity-separable hypothesis depicted “shears” along the vertical (temporal frequency) direction, rather than the direction orthogonal to the preferred velocity plane. See Methods for details. (h-i) Contour plots of slices through the two selectivity volumes from (f) and (g) at the optimal direction, superimposed with stimuli for two “classical” tuning experiments (black lines) containing the optimal stimulus (black ball) and suboptimal stimuli (dark gray): (h) spatial frequency tuning at optimal and low temporal frequencies, and (i) temporal frequency tuning at optimal and low spatial frequencies. (j-k) Temporal and spatial frequency tuning for the two models is the same for “classical” stimuli (red-blue dashed lines), but different for non-optimal stimuli (red and blue solid lines). The velocity-separable (light red) spatial and temporal frequency tuning curves are shifted away from the tuning curves observed for optimal stimuli.

Speed tuning and pattern motion selectivity in MT were typically studied separately. Furthermore, previous studies in MT were performed in at most two of three dimensions: spatial and temporal frequency (Perrone and Thiele, 2001; Priebe et al., 2003; Priebe et al., 2006), or direction and speed (Rodman and Albright, 1987). Recently, Nishimoto and Gallant (2011) and Inagaki et al. (2016) quantified MT selectivity in all three dimensions simultaneously, but did not relate their findings to pattern motion selectivity.

The Simoncelli and Heeger (1998) model of MT motion computation proposes that speed tuning and pattern motion selectivity both emerge from selective weighting of V1 afferents, parameterized in all three frequency dimensions. The model posits that MT neurons sum responses of V1 neurons whose preferred stimuli are consistent with a common velocity. MT neurons could, however, sum V1 afferents whose preferences share a common temporal frequency.

Here, we unify previous theory and experimental data in a coherent framework, by modifying the Simoncelli and Heeger (1998) model to allow direct fitting to electrophysiological recordings. Specifically, we compared the two hypotheses of MT computation above in their ability to explain the responses of neurons in areas V1 and MT of anesthetized and awake macaques to a large collection of sinusoidal gratings and plaids (superimposed gratings with different orientations and temporal frequencies). We fit these responses with a linear-nonlinear model of MT computation, in which the MT receptive field was constructed by summing velocity-specific or temporal frequency-specific combinations of V1 afferents. We refer to the former model variant, in which selectivity to spatial and temporal frequency varies jointly, as the *velocity-separable* model, and the latter model as the *frequency-separable* model. Nearly all V1 neurons were better described by the frequency-separable model. When probed with drifting sinusoidal gratings, MT responses were equally well-described by both models. However, when probed with plaid stimuli, the velocity-separable model systematically outperformed the frequency-separable model for pattern-selective neurons. This is the first direct evidence establishing speed tuning and pattern motion selectivity in area MT as consequences of a single organizing principle: selectivity organized along a preferred velocity plane.

## Materials and Methods

### Anesthetized recording procedures

We recorded from 7 anesthetized, paralyzed, adult male macaque monkeys (*M. fascicularis*) and one adult female macaque (*M. mulatta*) using standard procedures for surgical preparation and single-unit recording, as described previously (Cavanaugh et al., 2002). We maintained anesthesia and paralysis by intravenously infusing sufentanil citrate (6-30 *µ*g kg*^−^*^1^ h*^−^*^1^), and vecuronium bromide (Norcuron, 0.1 mg kg*^−^*^1^ h*^−^*^1^), respectively, in isotonic dextrose-Normosol solution (4-10 mL kg*^−^*^1^ h*^−^*^1^). Vital signs (heart rate, lung pressure, electroencephalogram (EEG), electrocardiogram (ECG), body temperature, urine flow and osmolarity, and end-tidal CO_2_ partial pressure (pCO_2_)) were continuously monitored and maintained within appropriate physiological ranges. Atropine was applied topically to dilate the pupils. Gas-permeable contact lenses protected the eyes, which were refracted with supplementary lenses chosen by direct ophthalmoscopy. Experiments typically lasted 5-7 days at the end of which the monkey was killed with an overdose of sodium pentobarbital. We conducted all experiments in compliance with the US National Institutes of Health Guide for the Care and Use of Laboratory Animals and with the approval of the New York University Animal Welfare Committee.

The monkey was positioned so his eyes were 57-114 cm from the display. Grating and plaid stimuli each lasted for 1,000 ms and were presented in randomly interleaved blocks. We used 0.5-3MΩ impedance quartz-platinum-tungsten microelectrodes (Thomas Recording) to make extracellular recordings in the brain through a craniotomy and small durotomy. Electrolytic lesions were made at the end of each recording track for histological confirmation of MT recording sites. For each isolated unit, we determined eye dominance and occluded the non-preferred eye. While isolating neurons in V1 for recording, we selected those with strong direction tuning.

### Awake recording procedures

To verify that the observations we made in the anesthetized preparation were not affected by anesthesia, we also recorded from 2 awake, actively fixating, adult male macaques (one *M. mulatta* and one *M. nemestrina*). No differences were observed between awake and anesthetized data. A headpost was surgically implanted for head stabilization using the design and methods described in (Adams et al., 2007). In a second surgical procedure, a chamber was implanted for chronic electrode recording over the superior temporal sulcus (STS) of the left hemisphere, using the techniques and a variant of the design described in (Adams et al., 2011). Prior to surgery, we used structural MRI and Brainsight software (Rogue Research, Canada) to design a chamber with legs matched to the curvature of the monkey’s skull (Johnston et al., 2016) above the STS.

We acclimated each monkey to his recording chair and experimental surroundings. After this initial period, he was head-restrained and rewarded for looking at the fixation target with dilute juice or water. Meanwhile, we used an infrared eye tracker (EyeLink 1000; SR Research, Canada) to monitor eye position at 1000Hz via reflections of infrared light on the cornea and pupil. The monkey sat 57 cm from the display.

The monkey initiated a trial by fixating on a small white spot (diameter 0.1°), after which he was required to maintain fixation for a random time interval between 2,350 and 4,350 ms. A grating or plaid stimulus would appear 100 ms after fixation began and last for 250 ms. Stimulus conditions were presented in randomly interleaved blocks. The monkey was rewarded if he maintained fixation within 1-1.75° from the fixation point for the entire duration of the stimulus. No stimuli were presented during the 300 to 600 ms in which the reward was being delivered. If the monkey broke fixation prematurely, the trial was aborted, a timeout of 2,000 ms occurred, and no reward was given.

For extracellular recordings, we advanced 0.5-3MΩ impedance tungsten microelectrodes with epoxylite insulation (FHC, Bowdoin, ME) through a 23G stainless steel dura-penetrating guide tube. We identified area MT from gray matter-white matter transitions and isolated neurons’ brisk, direction-selective responses.

### Analysis of neuronal response

We used Expo software (http://corevision.cns.nyu.edu) on an Apple Mac Pro to process signals and sort single units. Signals were amplified 1000×, bandpass filtered (300Hz to 10kHz), fed into a multiple-window time-amplitude discriminator, and time-stamped with 100*μ*s resolution. During each single-unit isolation attempt, discriminator windows and thresholds were manually set, online, to most unambiguously and stably distinguish a single spike waveform from noise and other units. If spike waveforms from multiple neurons overlapped to the extent to which they could not be separated, the discriminator allowed us to detect this based on sub-refractory period interspike intervals. If this type of multiunit activity was detected, we would then either manually refine the discrimination windows to effectively isolate the unit, move the electrode a small distance, or abandon that neuron for experimentation and move on to the next unit that could be fully isolated. During the experiments, each unit waveform was continuously monitored for isolation stability. If necessary, minor, manual adjustments detailed above were made to preserve isolation quality. Spike waveform consistency and interspike interval distributions over the entire duration of the experiments were verified post hoc. While running experiments, Expo automatically generated and displayed tuning curves, from which half-maximum spiking response rates, and their corresponding stimulus values, could be directly computed and displayed through the graphical interface.

### Visual stimulation

We presented visual stimuli on a gamma-corrected CRT monitor (an Eizo T966 during anesthetized experiments, and an HP P1230 during awake experiments; mean luminance, 33 cd/m^2^) at a resolution of 1,280 *×* 960 with a refresh rate of 120 Hz. Stimuli were generated and presented on the same Mac Pro using Expo.

For each isolated unit, we presented windowed sinusoidal grating stimuli to determine, by hand, initial estimates of each cell’s receptive field and preferred size. We used a standard sequence of tuning experiments to make precise estimates of the cell’s tuning preferences. Each tuning curve measured in this sequence featured 100% contrast single gratings varying along a single stimulus dimension, beginning with size tuning and followed by direction, spatial frequency, and temporal frequency tuning. After each of these individual tuning experiments finished, we determined the preferred stimulus value of the dimension tested and used it in subsequent experiments. Next, we measured pattern direction selectivity, at optimal spatial and temporal frequencies, with interleaved drifting gratings and plaids. In the rare cases in which this experiment yielded a different grating direction preference from the previously determined value, we repeated the full sequence of tuning curve measurements, to make sure optimal values would be used for the single component and planar plaid experiments which followed. The receptive fields of all recorded neurons were centered between 2° and 30° from the fovea.

Next, we ran the planar plaid study, which required no further stimulus optimization. In the planar plaid study, stimuli were chosen to span four different direction tuning curves at the optimal spatial frequency (see figure 7(a,b)). The first two were based on single gratings at 50% contrast, one with temporal frequency held constant at the optimal value (“constant frequency”), presented in all directions in 30° intervals, and the other with constant, optimal velocity (“constant velocity”) from -90°to 90° relative to the preferred direction, in 15°intervals. Since a given velocity corresponds to spectral content lying on a tilted plane in frequency space, constant velocity gratings had a temporal frequency that varied with the cosine of their direction.

The last two tuning curves consisted of 120° “plaids” (sums of two gratings with orientations 120° apart). The component gratings had the same temporal frequency, and were presented at 50% contrast each. The “pattern direction” of motion (direction consistent with rigid translation, equal to the average direction of the two gratings) was sampled the same set of directions used for single grating tuning curves.

Following the planar plaid study, we ran the single component study. It included presentation of 225 drifting grating stimuli, each at 100% contrast. Stimuli were arranged to widely sample the three dimensions of spatiotemporal frequencies near a given neuron’s tuning preferences, using multiple tuning curves, each varying along a single dimension: direction, spatial frequency, and temporal frequency. These tuning curves were measured at optimal and suboptimal values, the latter of which were determined by reading out (or if necessary, linearly interpolating) the stimulus values which elicited a response at half the neuron’s maximum spike rate (computed directly in Expo, see Analysis of Neural Response above) in the preceding standard tuning curve experiments (see extended data table 4-1 for details). By sampling spatiotemporal frequencies in this way, we could efficiently concentrate stimuli to reveal subtle changes of each neuron’s selectivity in a manner that does not assume a particular shape of selectivity or manner of tuning specific to either V1 or MT.

Note that the first two direction tuning curves of the single component study differ from the two grating tuning curves in the planar plaid study in that in the planar plaid study: (1) gratings were at 50% contrast instead of 100%, and (2) constant frequency gratings spanned the whole range of directions rather than just the semicircle of directions centered at the preferred one. Even though gratings were only presented at 50% contrast in the planar plaid study, its implications for 3D frequency selectivity shape should generalize to the 100% contrast case, since differences in response strength for these contrast levels are negligible (Carandini et al., 1997; Sclar et al., 1990).

### Frequency- and velocity-separable models

The MT linear weighting functions for both the frequency- and velocity-based models are defined as a separable product of tuning functions over direction *w_d_*(*d*), spatial frequency *w_s_*(*s*), and temporal frequency *w_t_*(*t*).

Specifically, the frequency-separable linear weighting for a grating is defined as follows:

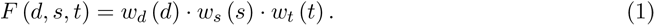

Direction tuning above is represented by a von Mises function:

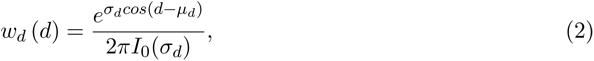

where *µ_d_* and *σ_d_* represent the direction preference and bandwidth, respectively, and *I*_0_() is the modified Bessel function of order 0 (which normalizes the integral of the numerator). Spatial frequency is represented by a logNormal function, parameterized by spatial frequency preference *µ_s_* and bandwidth *σ_s_*:

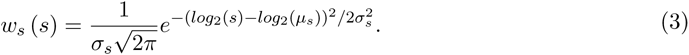

Finally, temporal frequency is represented by a Gaussian in coordinates which are linear at low frequencies and logarithmic at higher ones:

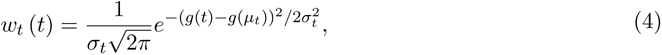

where *g*(*t*) is

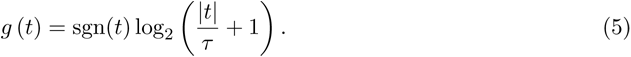

Using this functional form for temporal frequency tuning allows *w_t_* (*t*) to be logarithmic at high temporal frequencies, but also be zero-valued and continuous at zero temporal frequency. The parameter *τ* determines the temporal frequency at which the function transitions from linear to logarithmic, and *µ_t_*and *σ_t_* are the temporal frequency preference and bandwidth, respectively.

The velocity-separable linear weighting function is defined as follows:

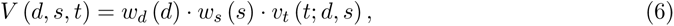

where the velocity-separable temporal frequency function, *v_t_*, is defined as a Gaussian, again linear at low frequencies and logarithmic at higher ones:

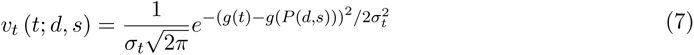

The only difference between *w_t_*(*t*) (equation (4)) and *v_t_*(*t*; *d, s*) (equation (7)) is that in the latter, temporal frequency tuning is centered on the preferred speed plane *P* (*d, s*):

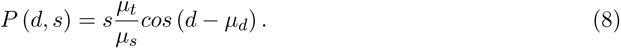

Note that a consequence of equations (6)-(8) is that the velocity-separable model “shears” vertically in the direction of the temporal frequency axis, rather than in the direction orthogonal to the preferred velocity plane (see figure 1). This was done deliberately, to account for the broad temporal frequency tuning MT neurons exhibit near the preferred direction and spatial frequency (see figures 4 and 8).

Previous models included a V1 normalization stage, either explicitly or implicitly simulated, at this part of the computation (Simoncelli and Heeger, 1998; Rust et al., 2006; Nishimoto and Gallant, 2011). Normalization at the V1 stage has been previously shown to be an important contributor to MT tuning properties. While we could have made it an explicit piece of the model here, Rust et al. (2006) showed that, in some cases, it can be combined with the MT normalization stage to yield a single normalization computation. Moreover, V1 contrast normalization is engaged only when contrast varies widely, and cross-orientation suppression is strongest for components close in orientation. Since the grating components in the plaid stimuli in our experiments are always 50% contrast and 120° apart, we assume such cross-orientation and contrast normalization effects in V1 are negligible. Thus, we assume the next model stage consists of MT neurons summing the responses to each plaid component.

Finally, the full MT model response is computed by raising the linear responses to a power *β*, and then normalizing them:

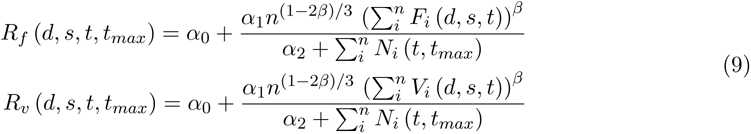

where the sums are over the components of the stimulus (*n* = 2 for plaids, *n* = 1 for gratings), and the *α*_0_ and *α*_1_ parameters represent the spontaneous and maximum discharge rates of the cell. The relative gains of responses to grating and plaid are controlled by the *n*^(1*−*2*β*)*/*3^ term in the numerator in equation (9).

The normalization signal, *N_i_*(*t, t_max_*), is meant to approximate the effects of tuned normalization. In the original Simoncelli and Heeger cascade model, MT normalization signals were computed by summing over a simulated population of MT neurons, but this construction would be computationally prohibitive in the context of fitting the model to spiking data. We parameterize the tuning as follows:

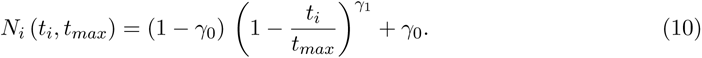

This function is maximally active at zero temporal frequency (with a value of 1) and minimally active with a value of *γ*_0_ at *t_max_*. *t_max_* is the highest temporal frequency simulated and experimentally presented. We used this form of normalization for fitting tractability and because it has useful properties, namely, it: (1) ensures there is no suppression at the preferred temporal frequency, (2) can be completely disabled by setting *γ*_0_ = 1, and (3) can be sub-linear, linear, or super-linear.

The rationale for this particular approximation of MT tuned normalization is based on the following thought experiment. We start with the assumption that neurons in the MT normalization pool fall into component-, intermediate-, and pattern-selective subpopulations (see figure 1(c)). Next, we assume all neurons’ selectivities in each of these subpopulations evenly tile the space of spatiotemporal frequencies. Now we will consider the consequences of summing together the spatiotemporal frequency selectivities of all the neurons in each subpopulation. Specifically, the manner in which these selectivities sum together and overlap each other in frequency space determine whether and how normalization is tuned. Any systematic biases in tuning overlap, as a function of spatiotemporal frequency, yield tuned normalization.

Component neurons from both model variants have narrow tuning, overlapping only between neurons with adjacent spatiotemporal tuning preferences. As a consequence, the summed responses of the population of component neurons evenly tile frequency space, producing an untuned normalization signal.

Frequency-separable pattern-selective neurons have broad direction tuning, so their overlap will occur most strongly in direction. The overlap, however, will be separable in spatial and temporal frequency, so for any subpopulation with the same spatial and temporal frequency tuning at all directions, the overlap will be confined to a donut-shaped region centered on those spatial and temporal frequencies. Since we assume the population of frequency-separable pattern selective neurons are evenly distributed across all preferred spatial and temporal frequencies, the tuning overlap will also be evenly distributed, yielding an untuned normalization signal.

Finally, velocity-separable pattern-selective neurons, which are organized along tilted planes which pass through the origin, will have strong overlap at zero and low temporal frequencies regardless of their preferred direction. As such, a pool of velocity-separable pattern neurons tiling frequency space generate a tuned normalization signal which strongly emphasizes low/zero temporal frequency. The same conclusions can be drawn for intermediate neurons in each separable model, although their selectivities will overlap less, yielding a similar, but weaker, tuned normalization signal.

### Estimating model parameters for individual cells

In total, the model has 9 free parameters for the single-grating study and 10 for the planar plaid study. For the former, they are: the direction preference and bandwidth (*µ_d_* and *σ_d_*), spatial frequency preference and bandwidth (*µ_s_* and *σ_s_*), temporal frequency preference, bandwidth, and log-linear transition (*µ_t_*, *σ_t_*, and *τ*), and the spontaneous and maximum firing rates (*α*_0_ and *α*_1_). For the latter experiment, *µ_s_*, *σ_s_*, and *µ_t_* are unconstrained by the data and are therefore held fixed at experimentally determined values, but the exponent (*β*), semi-saturation constant (*α*_2_), and normalization parameters (*γ*_0_ and *γ*_1_) are free. To avoid model fits producing spuriously wide temporal frequency tuning, we included temporal frequency tuning data collected immediately prior in the fitting of the planar plaid dataset. That temporal frequency tuning data, along with the planar plaid stimuli which sample different directions, constrain *µ_d_*, *σ_d_*, and *σ_t_*. In each study, the frequency- and velocity-separable models have the same parameters, and only differ in the parameterization of their temporal frequency linear weighting functions, *w_t_* and *v_t_* (see equations (4) and (7)).

For each cell, we optimized the model parameters by minimizing the negative log-likelihood (NLL) over the observed data, assuming spike counts arise from a modulated Poisson model. An additional parameter, *σ_G_*, describes across-trial fluctuations in neural response gain (Goris et al., 2014) and was optimized to the data independently from the frequency- and velocity-separable models and held constant during model fitting. We performed the optimization in successive steps, using optimal values from one step as initialization values for the next. First, we fit *τ*, then added the rest of the MT linear weighting parameters, and then in the case of the planar plaid experiment, the MT parameters controlling the MT nonlinearity.

### Experimental design and statistical analysis

In the single component study, we recorded single-unit responses of 13 V1 neurons and 39 MT neurons from seven anesthetized, paralyzed, adult male macaque monkeys (*M. fascicularis*) and one adult female macaque (*M. mulatta*). From those same eight monkeys, we recorded 21 V1 neurons and 53 MT neurons for the planar plaid study. These monkeys were also used for other visual system experiments not reported here—12 of the 53 MT neurons reported had another experiment run after the single component and planar plaid studies were run. Of the 53 MT neurons in the planar plaid study, 23 were also in the single component study. All of the 13 V1 neurons from the single component study are in the set of 21 V1 neurons in the planar plaid study. We additionally recorded 58 MT neurons from two awake, actively fixating, adult male macaques for the planar plaid study (31 from *M. mulatta* “Albert” and 28 from *M. nemestrina* “LW”). In all studies, neurons were only excluded from analysis if spike isolation degraded during the experiment, or if spike rates were too low (e.g., always below 10 spikes/sec) or variable to reliably predict direction tuning. No additional neurons were rejected from any subsequent analyses.

Following stimulus onset, we counted spikes within a 1,000 ms window (anesthetized experiments) or a 250 ms window (awake experiments). For each cell, latency of these windows (relative to stimulus onset) were chosen by maximizing the sum of response variances computed for each stimulus condition (Smith et al., 2005). Error bars on tuning curve responses indicate *±*1 standard deviation.

We used standard methods to compute each cell’s “pattern index” (Movshon et al., 1985; Smith et al., 2005). First, we computed partial correlations between the actual (constant temporal frequency) plaid responses and idealized predictions of pattern and component direction selectivity (*r_p_* and *r_c_*, respectively). We then converted these values to Z-scores to stabilize the variances of the correlations (*Z_p_* and *Z_c_*). Finally, the pattern index is the difference of these two quantities: *Z_p_ −Z_c_*. Cells were classified as pattern selective if *Z_p_ −Z_c_ >* 1.28, or component-selective if *Z_c_ −Z_p_ >* 1.28. Both thresholds correspond to a significance of *P* = 0.90. Confidence intervals on pattern index were computed from the 95th percentile of 100 bootstrapped estimates (Efron and Tibshirani, 1993; Rust et al., 2006).

For optimal and non-optimal spatial and temporal frequency tuning curves in the single component study in figure 2(b-d), we fit a difference of log2-Gaussians (Hawken et al., 1996). For each neuron, the stimulus value corresponding to the peak of this fitted difference of log2-Gaussians function was used as the fitted preferred stimulus in the figure. To test the robustness of these tuning curve fits, we ran a bootstrap analysis in which trials from each tuning curve were pseudo-randomly resampled 1000 times, with replacement, with the restriction that no stimulus condition had zero trials sampled. The error bars in figure 2(b-d) represent the 95% confidence intervals of these bootstrapped fitted peak stimulus values. Some tuning curves had flat tops, yielding unreliable tuning preference estimates. We therefore excluded neurons (7 MT, 0 V1) from all analyses in figure 2(b-d) which had a confidence interval exceeding 1.5 decades in any of the three. The conclusions are the same with or without these neurons.

**Figure 2.**
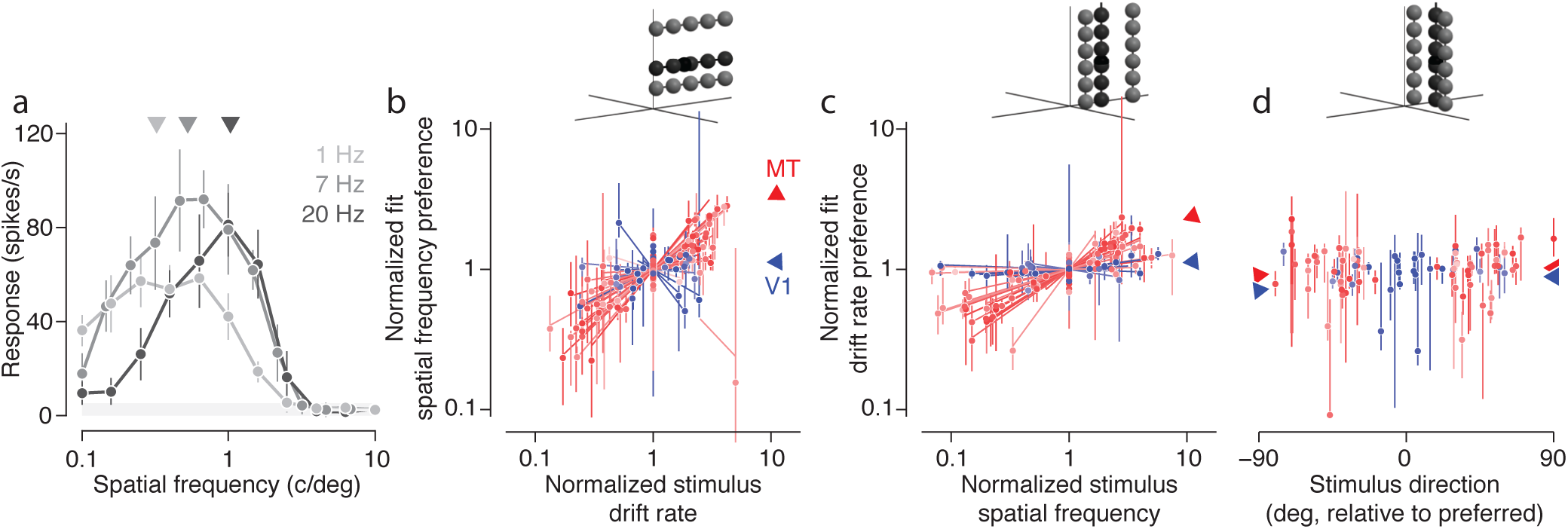
At the preferred direction, V1 is frequency-separable and MT is velocity-separable. (a) Spatial frequency tuning curve data from an example MT cell, measured at three temporal frequencies (error bars denote *±*1 s.d.). The light gray shaded area denotes the spontaneous firing rate, *±*1 s.d. The fitted SF tuning preferences for the three curves are shown above as triangles. (b) The fitted SF preferences from each cell are plotted against the TFs at which they were presented. Both axes are on a normalized scale, representing the ratio of the non-optimal frequencies, relative to the optimal frequency. Each line is the best fit line to the data for one cell. The data along the ordinate axis are aligned to the offset of each best fit line. Lines and points are shaded by the pattern index corresponding to each individual cell. Red corresponds to MT neurons, with darker shades corresponding to higher pattern index, and blue corresponds to V1 neurons, with darker shades corresponding to lower pattern index. The blue and red triangles indicate the mean slopes for all V1 and MT neurons, respectively. Error bars indicate the 95% confidence intervals of 1000 bootstrapped fitted peak stimulus values (see Methods for details). (c) Same as (b), but based on TF tuning curves measured at optimal and suboptimal SFs. (d) Same as (c), but based on TF tuning curves at optimal and suboptimal directions. Here the points are aligned to the origin, which represents the preferred TF at the preferred direction. Mean slopes in (d) are computed separately for the different suboptimal directions.

For the constrained parameter search during model fitting, we used a simplex algorithm (the Matlab function ‘fmincon’). To avoid overfitting and obtain estimates of parameter stability (i.e., the error bars in figures 9(a,b) and 10), we fit the model on 100 bootstrap resamplings of the data. Bootstrapping was done on a per stimulus-condition basis—that is, trials within each stimulus condition were sampled with replacement, ensuring that there were no stimulus conditions without data. Error bars on model fits indicate *±*1 standard error.

To compare model fits to a given neuron, we computed “velocity superiority”, the difference of the normalized NLLs of the velocity- and frequency-separable models. The NLLs were normalized by their corresponding “null” and “oracle” models, which serve as lower and upper bounds, at 0 and 1, respectively. The null model assumes the cell has two possible response rates: one when a stimulus is present and another when there is no stimulus. These are fixed to the measured mean spontaneous and maximal stimulus-driven response rates, respectively. The oracle model serves as an upper bound for the models’ performance, computed by using the measured mean responses to each stimulus condition to predict the neuron’s response to any individual presentation of that stimulus. We used the Wilcoxon signed rank test to test velocity superiority significance and Pearson’s *r* to assess correlation between velocity superiority and other quantities, such as pattern index.

## Results

### Joint and independent representations of motion in the frequency domain

Any image sequence can be decomposed (using a three-dimensional Fourier transform) into a sum of sinusoidal gratings of differing orientation, and spatial and temporal frequency. A single point in this 3D frequency domain corresponds to a drifting sinusoidal grating with a unique orientation, spatial frequency, and temporal frequency (figure 1(d)). More complex spatial patterns contain mixtures of gratings of different orientations and spatial frequencies. If these patterns are rigidly translating over time, their frequency domain constituents lie on a tilted plane through the origin, the slope of which is equal to the object’s speed (figure 1(e); Watson and Ahumada (1983); Watson and Ahumada (1985)).

How do V1 and MT neurons represent visual motion? Most V1 neurons are selective for a relatively narrow range of orientations, and spatial and temporal frequencies, corresponding to a ball in the frequency domain (Goris et al., 2015). If MT neurons are specialized for analyzing rigid motion, their receptive fields should be organized along just such a plane with slope equal to a preferred speed (figure 1(f), “velocity-separable”) (Simoncelli and Heeger, 1998). While there is some direct physiological evidence for velocity-separable organization (Rodman and Albright, 1987; Perrone and Thiele, 2001; Priebe et al., 2003; Nishimoto and Gallant, 2011), as well as perceptual evidence (Adelson and Movshon, 1982; Schrater et al., 2000), this is not the only kind of receptive field organization consistent with known MT properties.

However, almost all experimental measurements of grating direction selectivity use stimuli that lie along a horizontal plane of constant temporal frequency. By treating spatial and temporal frequency independently, they implicitly assume that MT direction selectivity is organized along these planes (“frequency-separable,” figure 1(g)). Evidence exists for this alternative possibility (Perrone and Thiele, 2001; Priebe et al., 2003)—an MT neuron with this type of organization would still be direction-selective, but in a manner that is more strongly influenced by variations in spatial pattern.

These two model structures make different, testable predictions about how MT tuning should change in response to preferred and non-preferred stimuli. Experimenters typically assess a neuron’s spatiotemporal frequency tuning preferences by presenting gratings varying along one of the three dimensions of the frequency domain, while keeping the values in the other two dimensions fixed at the best estimate of the neuron’s preferences (black lines in figure 1(h-i) for spatial frequency, temporal frequency, and direction, respectively). For a simulated neuron, the tuning curves (red and blue dashed lines in figure 1(j-k)) generated from these optimized stimuli have their peaks at the neuron’s preferred spatiotemporal frequency (represented as the black points in figure 1(h-i)). The two predictions differ most for tuning curves measured at non-preferred frequencies (figure 1(h-i), dark gray lines and points), most notably in the tuning curves’ peak locations. The frequency-separable hypothesis predicts tuning in response to stimuli of non-preferred spatial and temporal frequency that is lower in amplitude but with a peak at the same frequency (blue lines, figure 1(j-k)). However, the velocity-separable hypothesis predicts that the tuning curve will shift (red lines, figure 1(j-k)), such that if the non-preferred tuning experiment is run at a frequency below preferred, the peak will also be at a lower frequency, and vice-versa.

To test these hypotheses in V1 and MT, we measured tuning curves at optimal and suboptimal spatial and temporal frequencies and asked whether or not there was a shift in their peak location. For “suboptimal” frequencies, we used the stimulus values corresponding to the half-maximum responses when measured at optimal frequencies (see Methods for details). Many cells (e.g., figure 2(a)), exhibited a peak spatial frequency tuning that increased with increases in grating temporal frequency, consistent with the velocity-separable hypothesis. To quantify this shift, and compare across neurons, we computed the peak spatial frequency and plotted it as a function of the relative temporal frequency at which it was measured (figure 2(b)). The degree to which the neuron is velocity tuned can be captured by the slope of the line through the data (0 for no speed tuning, 1 for ideal speed tuning, 0.37 for the neuron in (a)). V1 neurons show, on average, no slope (0.08 *±* 0.22 s.e.m., *n* = 13, blue in figure 2(b)), while MT neurons have a significantly positive slope (0.50 *±* 0.07 s.e.m., *n* = 39, red in figure 2(b)). Performing the same analysis for changes in temporal frequency preferences as a function of stimulus spatial frequency (figure 2(c)) yields similar slopes in MT (0.36 *±* 0.03 s.e.m.) and V1 (0.05 *±* 0.04 s.e.m.).

These measurements, all performed at the neuron’s preferred direction, support previous findings that V1 tends to be frequency-separable and MT velocity-separable (Simoncelli and Heeger, 1998; Perrone and Thiele, 2001; Priebe et al., 2003; Priebe et al., 2006; Nishimoto and Gallant, 2011). Since our goal was to characterize tuning in all three dimensions, we also assessed peak temporal frequency changes when measured at different directions (figure 2(d)), which should either remain constant or decrease (for the frequency- and velocity-separable hypotheses, respectively). When averaged across the populations, slopes were flat (figure 2(d), V1 (blue triangles) mean*−*0.001 *±* 0.004 s.e.m.; MT (red triangles) mean *−*0.0004 *±* 0.0007 s.e.m.), however, on a neuron-by-neuron basis, tuning at non-preferred directions was inconsistent. To probe the three-dimensional selectivity more finely, we presented stimuli at many more spatiotemporal frequencies, and fit velocity- and frequency-separable models directly to the responses.

### The velocity- and frequency-separable models

To examine MT receptive field organization in the frequency domain, we fit two modified versions of the Simoncelli and Heeger (1998) model of MT direction selectivity to the responses of individual neurons. Both models have the same structure: two stages, each with a linear weighting followed by a point nonlinearity and normalization (figure 3). The first (V1) stage consists of narrowly-tuned direction-selective complex cells, simulated with a linear weighting of a narrow band of frequencies, followed by squaring. The second (MT) stage computes a weighted linear combination of its V1 inputs, followed by another point nonlinearity and normalization.

**Figure 3.**
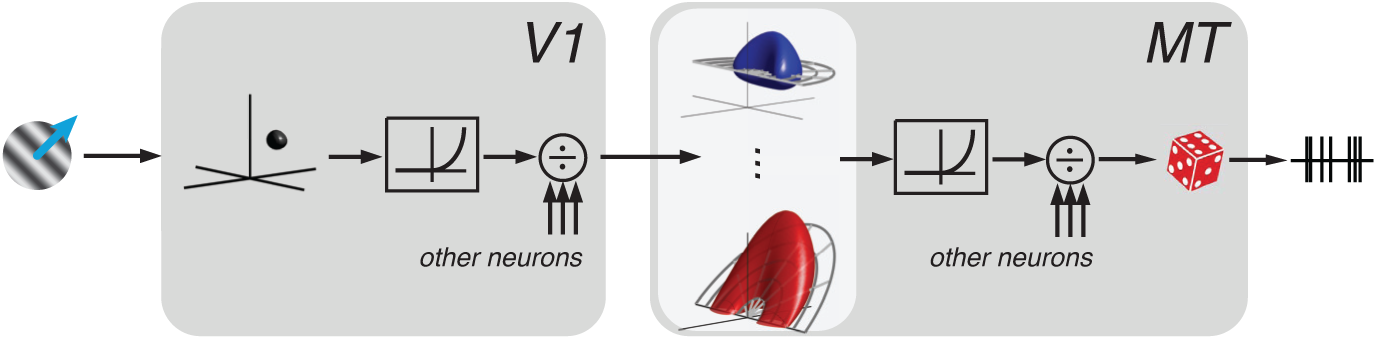
The separable models. A stimulus is passed through a narrowly tuned V1 linear weighting, then squared and normalized. V1 output is then passed to the MT neuron, which applies either a frequency- or velocity-separable linear weighting, then raises the output to a super-linear power, and undergoes another stage of normalization. Finally, a modulated Poisson process determines spike variability.

Linear weighting in the MT stage is the primary determinant of the MT neuron’s tuning properties, including pattern motion selectivity. We constrain it to be a separable product of three tuning curves. The first two (direction and spatial frequency tuning) are common to both models. In the frequency-separable model, the third separable function is temporal frequency tuning, independent of the other two dimensions. In the velocity-separable model, temporal frequency tuning co-varies with direction tuning such that the peak lies on a tilted plane whose slope is the preferred velocity of the neuron. This temporal frequency tuning parameterization is the only difference between the two models.

The MT stage nonlinearity controls interactions between multiple spatiotemporal frequencies simultaneously present in the stimulus, and thus plays an important role in establishing pattern motion responses. In the full models, the MT nonlinearity is composed of a point-wise power function, followed by divisive normalization. The divisive normalization operates on a uniform population of pattern and component cells which, taken in aggregate, are assumed to uniformly cover direction and spatial frequency, while exhibiting tuning for temporal frequency (see Methods for details). Single grating stimuli do not constrain this model component, and thus for the single grating study presented below, the exponent is fixed to a value of two.

### Single grating responses do not differentiate the models

How can we distinguish the two models? We designed a study in which we measured seventeen tuning curves, chosen on a neuron-by-neuron basis, to sample the frequency domain where the predictions of the models should deviate the most. The stimuli were full contrast sinusoidal gratings; five tuning experiments included a grating at the optimal spatiotemporal frequency, while twelve suboptimal tuning experiments did not (see methods and extended data figure 4-1 and table 4-1 for details). By comparing how responses fall off as stimuli deviate from the preferred spatiotemporal frequency, a picture of three-dimensional tuning should emerge in support of one model or the other.

**Figure 4.**
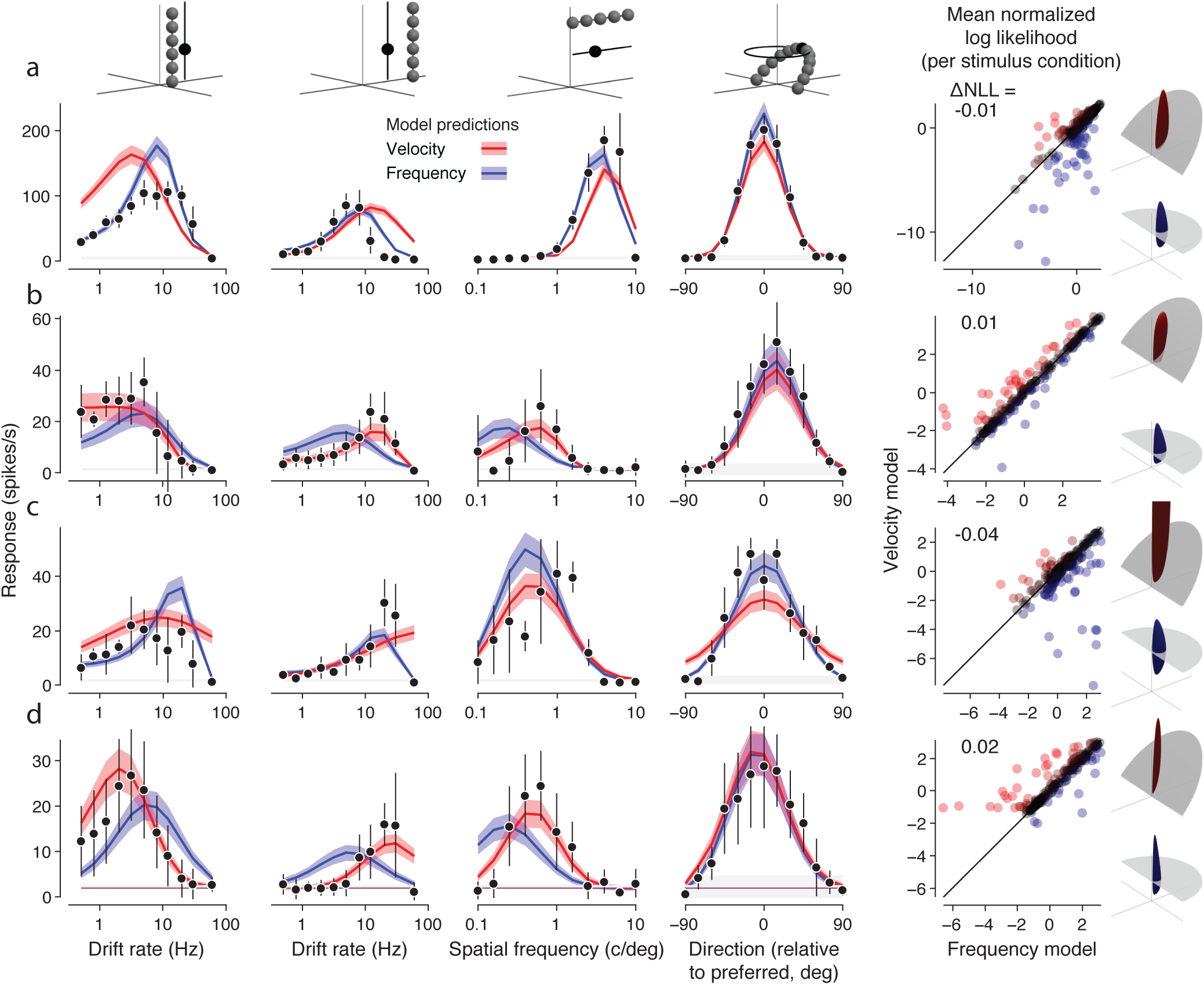
Comparison of actual and model-predicted responses to single gratings for four example MT neurons. (a,b) Two example component neurons, one better fit by the frequency-separable model (a) and one better fit by the velocity-separable model (b). (c,d) Two example pattern neurons, one better fit by the frequency model (c) and one better fit by the velocity model (d). Measured spike rate mean and standard deviation are shown in black. Velocity model predicted spike rates are shown in red, frequency model predictions in blue. All subsequent figures follow this color convention. Means are indicated by the dark lines, *±*1 standard deviation by the lighter shaded areas. In the scatter plots on the right, each point represents how well the frequency and velocity models predict the mean firing rate for one spatiotemporal frequency among the 225 presented across all experiments. Goodness of fit is expressed in terms of log likelihood under the modulated Poisson process, where values closer to zero indicate a better fit. The log likelihoods are normalized to a scale between 0 and 1, which represent the null and oracle model prediction log likelihoods, respectively (see Methods for details). Each point is colored on a Fisher transformed scale (i.e., in units of standard deviation). The difference between the velocity and model predictions for each neuron are summarized as a single value (Δ*NLL* = *NLL_V_ − NLL_F_*). Renderings of the frequency and velocity model linear weightings for each example neuron (rightmost column). All four neurons were recorded under anesthesia.

For each cell, we fit the frequency and velocity models to data from all 17 tuning experiments simultaneously. Figure 4(a-d) shows four of the seventeen tuning curves of the optimized model, fit to data from two example MT component neurons (figure 4(a,b)) and two MT pattern neurons (figure 4(c,d)). As expected, the models make substantially different predictions for spatial and temporal frequency tuning (first three columns in figure 4), but not direction tuning (fourth column). In the first two columns, for example, the velocity model predicts tuning peak shifts, whereas the frequency model does not.

Most tuning curves from each neuron are well fit by one of the two models (frequency model for figure 4(a,c) and velocity model for figure 4(b,d)), including changes in relative gain across tuning experiments. Relative model performance for each stimulus condition from all seventeen tuning experiments (points in the scatter plots in figure 4, rightmost column) show that while some spatiotemporal frequencies strongly distinguish the two models, most do not. This reflects the fact that some tuning curves are well-described by both models (e.g., the constant-velocity grating direction tuning, fourth column of figure 4).

This range of behavior was observed across the population. We assessed overall fit quality on a cell-by-cell basis by normalizing the log likelihoods of the models to null and oracle models. The null model assumes the cell has two possible response rates: one when a stimulus is present and another when there is no stimulus. These are fixed to the measured mean spontaneous and maximal stimulus-driven response rates, respectively. The oracle model serves as an upper bound for the models’ performance, computed by using the measured mean responses to each stimulus condition to predict the neuron’s response to any individual presentation of that stimulus. “Velocity superiority” is the difference of the normalized log likelihoods of the velocity-separable model and the frequency-separable model (figure 5).

**Figure 5.**
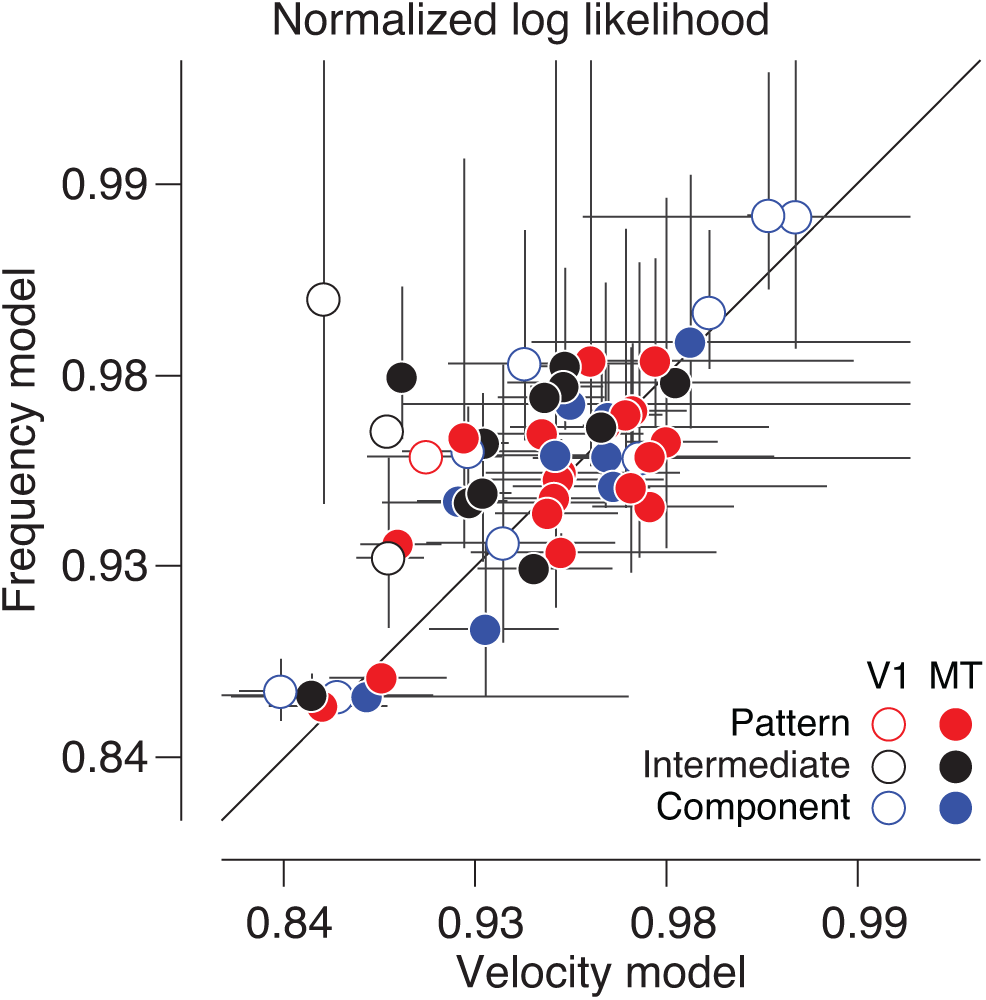
Single grating stimuli do not distinguish the two models. Fit quality, expressed as the normalized log likelihood of the velocity and frequency models, is plotted for each neuron on Fisher transformed axes. V1 neurons (*n* = 13) are shown with open circles, MT (*n* = 39) with closed circles. Blue, black, and red colors indicate whether a neuron is classed as component, intermediate, or pattern selective, respectively. Error bars denote standard error of the mean. On average, the two models are equally good at explaining the single grating MT data for any class of cells. The frequency model explains the V1 single grating data better.

In general, V1 cells were better fit by the frequency model (average velocity superiority of *−*0.03, *P* = 0.0046 Wilcoxon signed rank test, open circles in figure 5). Most MT neurons were clearly better fit by one model or the other, but overall, neither model was significantly better (*−*0.005 mean difference, *P* = 0.12 Wilcoxon signed rank test, filled circles in figure 5), regardless of pattern index.

Model fits to single grating responses provided further evidence that V1 neurons are frequency-separable, but were inconclusive for MT neurons. MT neurons tend to be velocity-separable for stimuli at the preferred direction (figure 2(b,c)) but have inconsistent tuning at off-directions (figure 2(d)). In theory, comparing direction tuning curves measured at either a given neuron’s optimal velocity or optimal temporal frequency should distinguish the models: they predict direction tuning bandwidth to be wider when the stimulus and model type match (e.g., velocity-separable model direction tuning for constant-velocity gratings should have a wider bandwidth than tuning for constant frequency gratings). In fact, measured tuning to these two stimuli are nearly identical for component neurons, and slightly broader, on average, for constant velocity gratings for intermediate and pattern neurons (figure 6). These data provide more evidence that MT neurons are likely velocity-separable, using different tuning measurements at non-preferred directions.

**Figure 6.**
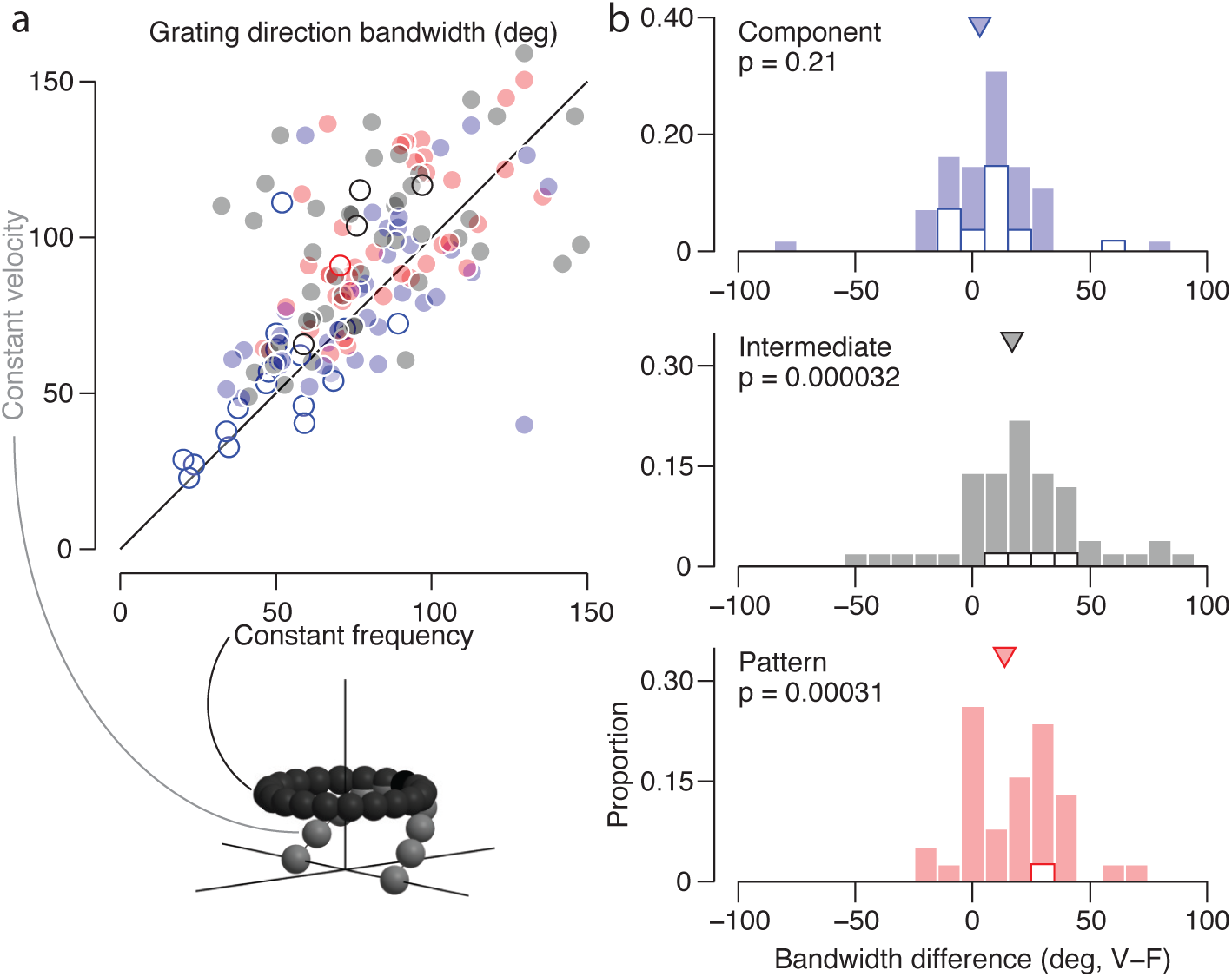
Velocity grating direction tuning tends to be slightly wider. (a) Direction bandwidth (in degrees) for each neuron was calculated separately for constant-frequency and velocity grating tuning curves. is plotted for each neuron. Blue indicates a component neuron, black intermediate, and red a pattern neuron. V1 neurons (*n* = 21) are shown with open circles, MT (*n* = 112) with closed circles. This figure includes isomorphic data recorded from the next experiment, described in the next section. (b) Differences of velocity and frequency grating bandwidth, by pattern classification. Proportions are expressed within each classification type, but including both V1 and MT neurons (open and filled stacked bars, respectively). Pattern and intermediate neurons have wider velocity grating direction bandwidth, significant below *p <* 0.0005 (Wilcoxon signed rank test).

Taking these observations into account, we concluded that single grating stimuli are not rich enough to fully distinguish the models. In particular, since only one spatiotemporal frequency is presented at a time, single gratings do not constrain the MT nonlinearity which underlies pattern computation. We therefore sought to use more complex stimuli and focused on sampling the frequency domain at non-preferred directions—those spatiotemporal frequencies which were the most informative in distinguishing the two models (both in theory and in practice).

### Compound stimuli reveal velocity-separable organization in MT

Selectivity for pattern motion is a defining property of MT neurons. Since single gratings alone are not rich enough to characterize MT, we ran a second study in which direction tuning curves were measured for gratings and 120° plaids, presented either at a given neuron’s optimal velocity or optimal temporal frequency (figure 7(a,b)). All stimuli were fixed at the neuron’s optimal spatial frequency. These stimuli can be equivalently described as gratings and plaids drifting either in the preferred direction of the cell or along the direction normal to the mean orientation (“constant velocity” or “constant frequency” in figures 7(a) and (b), respectively).

The two models make dramatically different predictions (figure 7(c)) for pattern-selective neurons: the frequency model predicts tuning for constant-velocity plaids to have a trough at the preferred pattern motion direction, and peak 90° from preferred. The velocity model predicts a near-constant, elevated response to all constant-velocity plaids.

**Figure 7.**
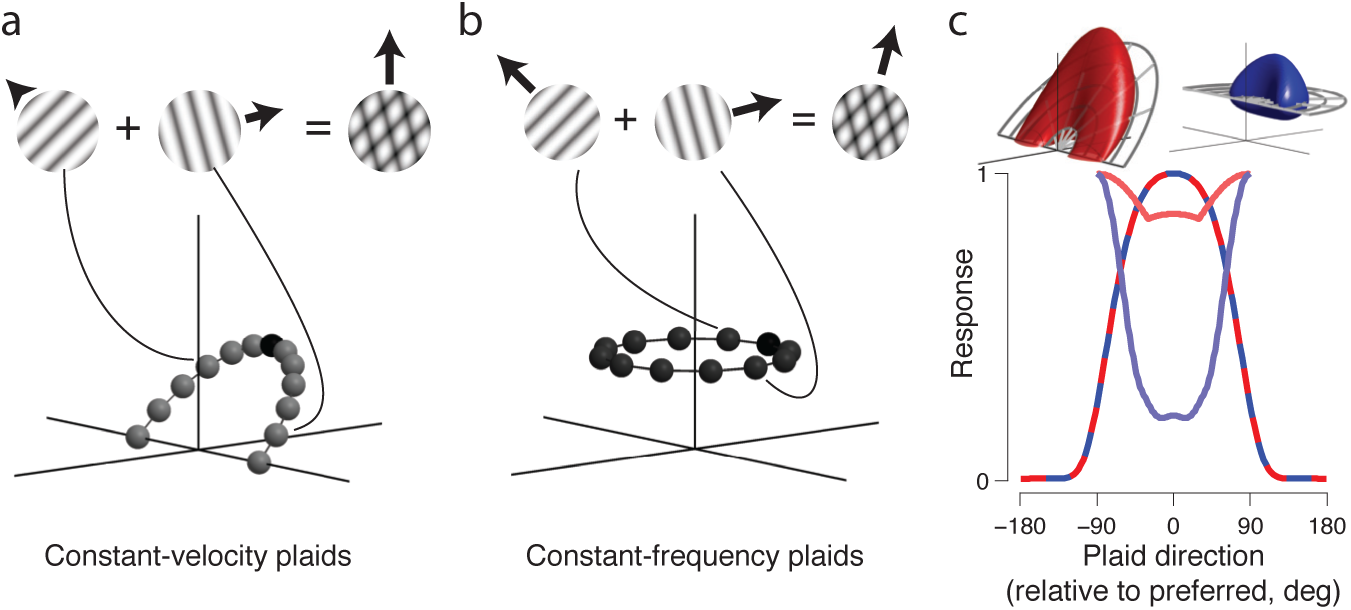
Two-component “planar plaid” experiment design and predictions. Constant-velocity and constant-frequency direction tuning experiments were done with gratings and plaids. Constant-velocity plaids (a) were constructed by superimposing two gratings 120° apart and drifting at a temporal frequency determined by the optimal velocity plane. Constant-frequency plaids (b) were two gratings 120°apart superimposed and drifting at the optimal temporal frequency. The example plaids shown contain the same orientations, but have different component drift rates, and thus different perceived drift directions. (c) For the two models matched in constant-frequency plaid direction tuning (red and blue dashed line), the velocity model (red) predicts a high response rate to all constant-velocity plaids. The frequency model (blue) is more narrowly tuned.

Responses to this new stimulus family were complex enough to fully constrain the models’ MT nonlinearity (a power function and divisive normalization—see methods for details). There were three free parameters in the nonlinearity that were fixed in the previous study. Since the spatial frequency preference and bandwidth and temporal frequency preference parameters were unconstrained by this dataset, they were fixed to values determined in preceding tuning measurements. In total, there was one additional free parameter fit compared to the single-grating study.

Qualitatively, the models predict that direction tuning should be flatter when the coordinate system of the model matches that of the stimuli. Two features of the measured responses stand out. First, constant frequency and constant velocity direction tuning curves to gratings are again clearly indistinguishable for all cells (two leftmost columns of figure 8). Second, the pattern selective MT neuron (figure 8(d)) exhibits much wider direction tuning bandwidth for constant velocity plaids as opposed to constant frequency plaids (fourth and third columns from the left in figure 8), while the other cells show more similar tuning bandwidth for the two plaid types. For all cells, both models capture grating responses well. However, the frequency model cannot account for the pattern selective neuron’s responses to both types of plaids simultaneously (figure 8(d), blue). The best it can do is pick a compromise direction tuning bandwidth that is too wide for constant frequency plaids and too narrow for constant velocity plaids. The velocity model, on the other hand, is able to account for all the data simultaneously, including the different plaid tuning bandwidths. This pattern cell is the only one of the four example cells that has substantial differences in the frequency- and velocity-separable model predictions. The increasingly divergent model predictions as pattern index increases is further illustrated by the increasingly different MT linear weighting functions (rendered in the last column of figure 8). The models make nearly identical predictions for narrowly tuned neurons, yielding velocity superiority indices at or near 0 (scatter plots on right of figure 8).

**Figure 8.**
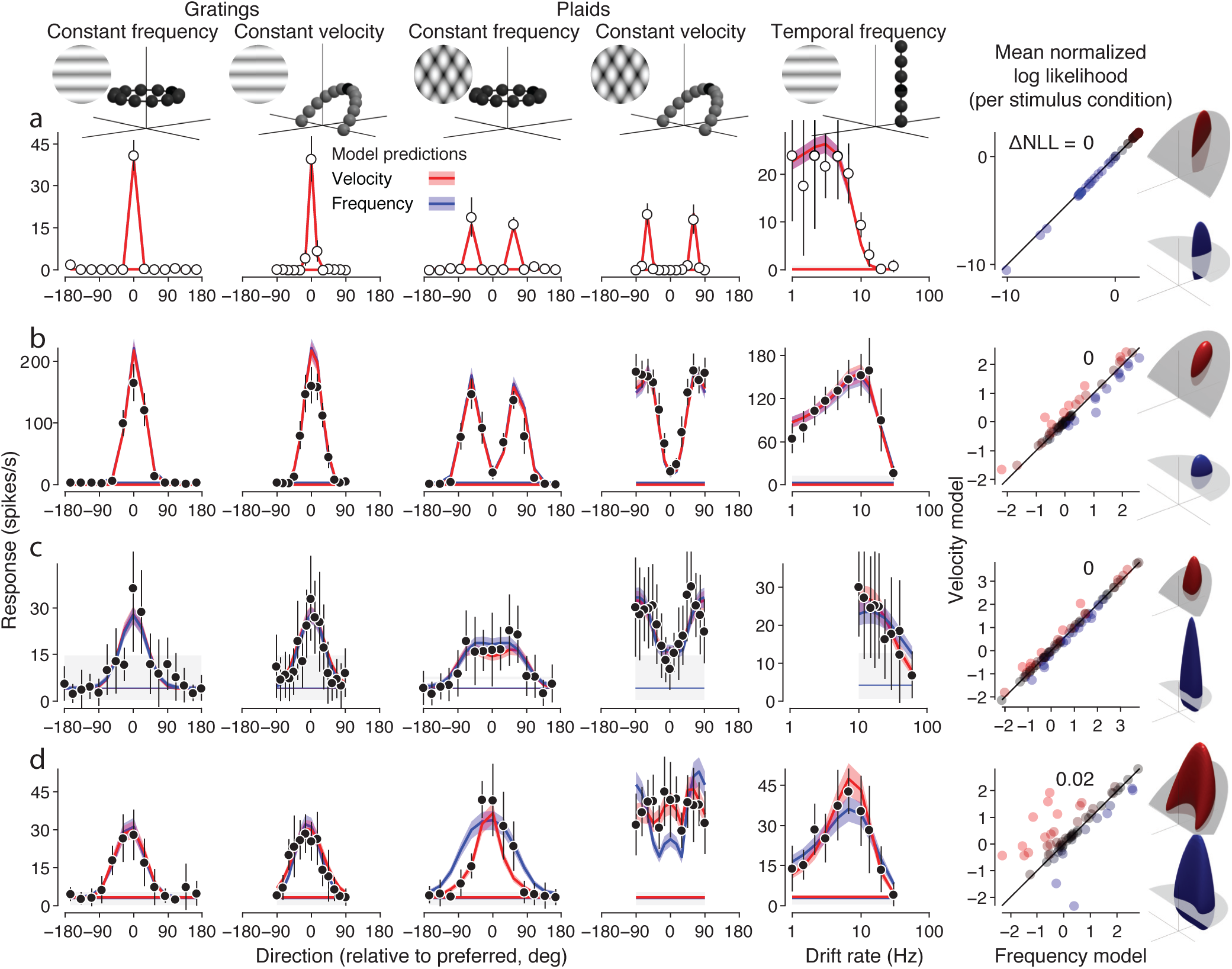
Comparison of actual and model-predicted responses to gratings and plaids for four example neurons. First five columns show data (points) and tuning curves predicted by the frequency-(blue) and velocity-separable (red) models. The first four columns are responses to gratings and plaids with constant frequency and velocity. The fifth column is temporal frequency data collected in a separate session, but included in the model fits. (a) a V1 component-selective neuron, (b) an MT component neuron, (c) an MT intermediate neuron, and (d) an MT pattern-selective neuron. The fifth column shows goodness-of-fit across all stimulus conditions, next to renderings of the fitted models. See figure 4 caption for details. Differences between the two model predictions become more apparent with increasing pattern selectivity. Neurons (a-c) are from recordings done under anesthesia, (d) is from an awake recording.

The relationship between velocity superiority and pattern index holds across cell populations. For the single grating data set, there is overall no significant correlation (figure 9(a), for all cells (Pearson’s *r* = 0.05, *P* = 0.70) or MT alone (*r* = −0.01, *P* = 0.95). There was, however, a significant negative correlation for V1 (*r* = −0.70, *P* = 0.007). Furthermore, there was no significant relationship between pattern index and the number of tuning curves per cell better fit by one model or the other (Pearson’s *r* = −0.22, *P* = 0.13). In contrast, responses to plaid stimuli indicate a significant correlation between velocity superiority and pattern index, and (figure 9(b), Pearson’s *r* = 0.34, *P* = 5.6*e* − 5). 86% of all pattern cells were better fit by the velocity model (*P* = 1.4*e* − 4, Wilcoxon signed rank test).

**Figure 9.**
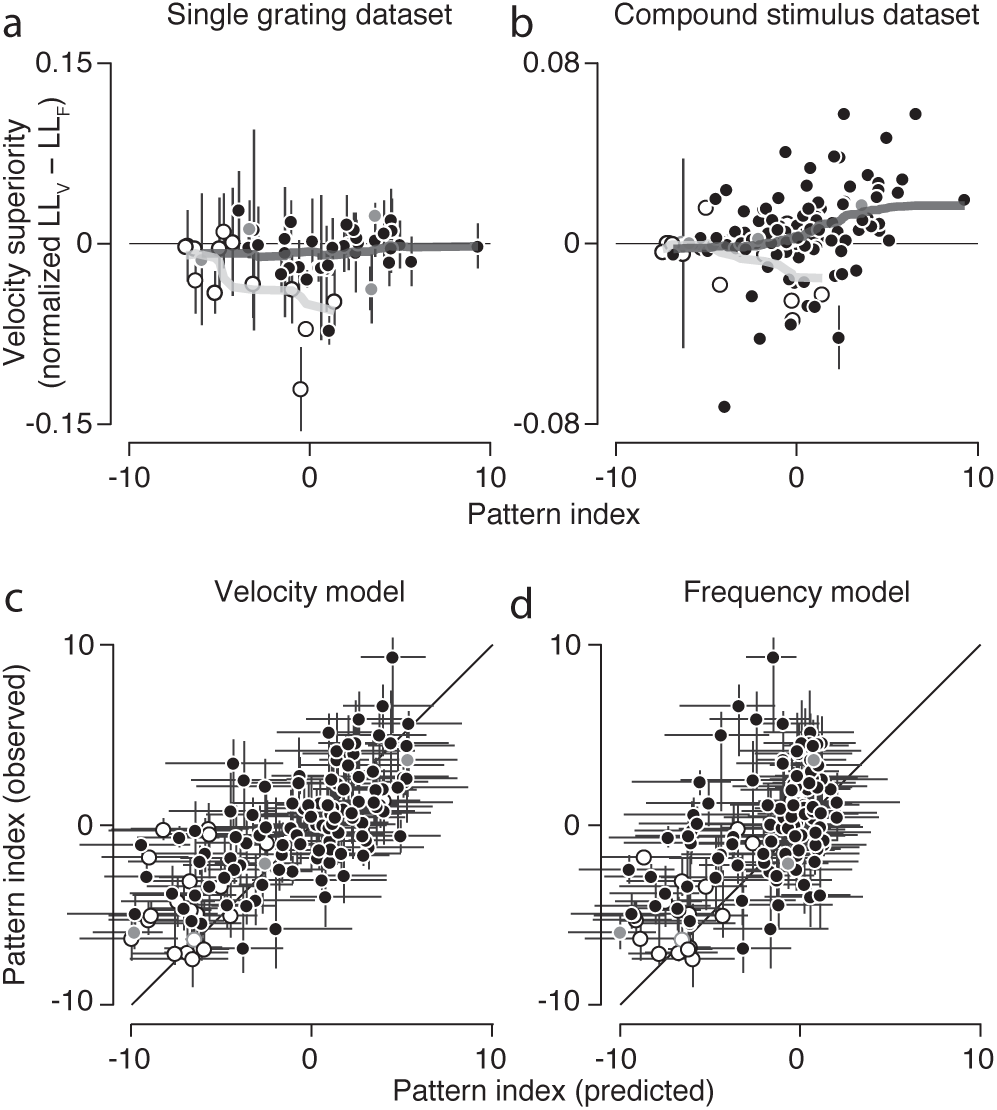
Compound stimuli reveal velocity-separable organization for pattern cells. (a,b) Velocity superiority, or the difference of normalized log likelihoods between the velocity and frequency models, per cell as a function of pattern index. V1 cells appear as open circles, MT closed. Example cells featured in figures 4 and 8 are highlighted in gray. Light and dark lines indicate the running mean, with a window of *±*1*/*3 of cells in each population. Error bars indicate *±*1 standard deviation, calculated from model fits to bootstrapped data (note most are smaller than the plotted points). (a) On average, for the single grating dataset, neither model better explains the single grating MT data (a) for any class of cells (*n* = 39). The frequency model explains the V1 single grating data better (*n* = 13). (b) Pattern cell responses to the compound stimulus dataset (V1: *n* = 21, MT: *n* = 112) are clearly better explained by the velocity model. Error bars indicate *±*1 standard error. (c,d) Observed and predicted pattern indices for each cell, derived from the compound stimulus dataset, for the velocity model (c) and frequency model (d). The velocity model can account for pattern index across all cell types, whereas the frequency model fails to predict the pattern selectivity of neurons classified as pattern-selective based on measured responses. Error bars indicate 95th percentiles, generated from pattern indices calculated by bootstrapping measured and predicted spike trains.

As a more direct test, we compared pattern selectivity of model predictions against the measured pattern selectivity of the cells. The velocity model accounts for the full range of pattern selectivity across the population (figure 9(c), Pearson’s *r* = 0.60). The frequency model, however, fails to produce any cells with pattern tuning (figure 9(d), Pearson’s *r* = 0.50), due to the compromises it must make when fitting both constant frequency and constant velocity plaid responses simultaneously.

### Model validation

Which characteristics are needed to describe the motion selectivity of a given neuron in MT? They are, in order of increasing complexity: (1) its preferred direction and speed, (2) the degree to which responses fall off as stimuli deviate from the preferred stimulus, and (3) the extent to which multiple overlapping motion components are treated independently or as a single, coherently moving pattern.

Experimentally, these attributes are established by identifying the stimulus that evokes the neuron’s maximum response, the shape of tuning curves for direction, spatial frequency, and temporal frequency, and calculating the pattern index based on correlating (constant frequency) grating and plaid direction tuning. The recorded direction curves we report (figure 8), with identical bandwidths for constant frequency and velocity gratings (all cells) and wider bandwidth tuning to constant velocity plaids (pattern cells), provide a novel, fourth criterion for describing MT motion selectivity.

The velocity model accurately captures the first, second, and fourth attributes (direction tuning, response fall-off, and matched/unmatched frequency bandwidths) by accurately reproducing tuning curves (figures 4 and 8). We verified that the velocity model also accounts for the third attribute (pattern motion selectivity), and accounts for the full range of pattern selectivity across the population (figure 9(c)). The frequency model, in addition to failing on the second and fourth criteria (figure 8), also fails on the third by failing to predict any pattern motion selective cells (figure 9(d)).

How does the velocity model provide a full account of motion selectivity? Capturing selectivity in all three frequency dimensions accounts for the first two attributes(direction tuning and response fall-off). Pattern selectivity, the third attribute, is controlled by increasing direction tuning bandwidth (Pearson’s *r* = 0.73, figure 10(a)) and the exponent in the nonlinearity (Pearson’s *r* = 0.60, figure 10(b)). Divisive normalization, which allows the model to adjust the fourth attribute(matched/unmatched frequency bandwidths), does so by producing grating direction tuning curves with more similar bandwidths than would be predicted in its absence. The semi-saturation, or “uniform” divisive normalization parameter, is very weakly correlated with pattern index (on a log2 scale, Pearson’s *r* = *−*0.16, *P* = 0.07, figure 10(c), see methods for details). This means that for neurons with higher pattern index, the temporal-frequency dependent suppression is stronger.

**Figure 10.**
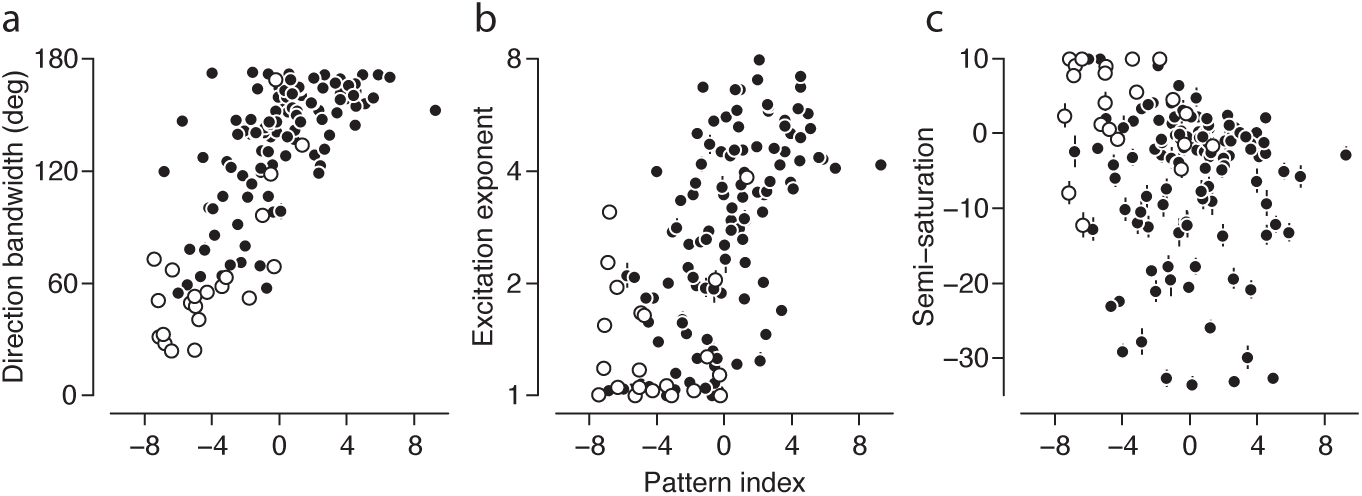
Relationship between velocity-separable model parameters and pattern index. Pattern index is strongly correlated with direction tuning bandwidth (a) and the log of the MT nonlinearity’s exponent (b). (c) Pattern index is negatively correlated with the log-2 of the semi-saturation, or “uniform” divisive normalization parameter. This means that for neurons with higher pattern index, the temporal-frequency dependent suppression is stronger.

Taken together, the two datasets and associated model fits reveal important aspects of MT computation. First, sinusoidal grating stimuli drifting in a neuron’s preferred direction can reveal a frequency-separable receptive field organization in V1, and a velocity-separable one in MT. These stimuli, however, are not sufficient to reveal the nonlinear behaviors that distinguish direction selectivity observed in MT from that observed in V1. Second, compound stimuli, which constrain nonlinear receptive field behavior in MT, reveal receptive fields that are organized along a neuron’s preferred velocity plane.

## Discussion

To date, attempts to characterize MT motion selectivity have generally followed two distinct strategies. They focused either on how multiple superimposed spatiotemporal frequencies are integrated into a single, coherent drifting pattern, or on how tuning varies across multiple dimensions of the spatiotemporal frequency domain. Here, we present a model that unifies these two approaches in a common framework, and for the first time, generalizes previous findings to all three dimensions of the spatiotemporal frequency domain.

We recorded responses of a large population of neurons in both MT and V1 to simple stimuli specifically designed to extensively quantify tuning in the spatiotemporal frequency domain as well as tuning for pattern motion. We fit two compact two-stage models to each individual neuron in the population. We found temporal frequency tuning in MT to be much broader than that instantiated in the Simoncelli and Heeger (1998) model. Furthermore, by comparing two models’ performance, we provide model-based evidence that MT neurons’ selectivity is best described by a tilted, constant velocity plane in the spatiotemporal domain. Finally, compound stimuli were necessary to reveal this organization—single sinusoidal gratings were not sufficient.

### Motion computation in the velocity-separable model

The separable models we developed and tested are generalizations of previous models (Simoncelli and Heeger, 1998; Rust et al., 2006). The Simoncelli and Heeger (1998) model was constructed using populations of V1 and MT neurons, each having their own rectifying nonlinearities and divisive normalization. The second (MT) stage of the model linearly weighted the afferent signals from V1 along a tilted, constant velocity plane in the frequency domain. But this model was not explicitly fit to single cell data, and comparisons of predicted to measured tuning curves were qualitative. The Rust et al. (2006) paper used a simplified model variant that predicted (and was fit to) responses to gratings and plaids at a single temporal frequency. The paper showed that pattern selectivity could be explained by incorporating opponent suppression and direction-tuned normalization. By fitting to a more diverse set of stimuli, and a model that includes a full range of temporal frequencies, we find that selectivity for both speed and direction of moving patterns can be captured in a single model. Note that we have incorporated temporal frequency dependent normalization in the MT stage (as opposed to the direction-tuned normalization of the Rust et al. (2006) model). For parameter values optimized to neurons in the compound stimulus dataset, this tends to sharpen direction tuning for constant velocity gratings.

In order to characterize MT receptive field structure in all three dimensions of the frequency domain, it was not feasible to simulate entire populations of V1 and MT neurons. By restricting our stimuli to gratings and plaids which would not be affected by normalization in V1, we could avoid explicitly simulating the V1 stage. Rather, the model evaluated tuning directly based on the separable product of tuning curves along three dimensions in the frequency domain. Since all three tuning curves are exponential functions, the separable tuning volume and exponent approximately accounts for both the linear weighting stages and power function nonlinearities of V1 and MT. As a result, three computational elements, all implemented in the MT stage, determine how a given MT neuron responds to moving stimuli: linear weights, a point-wise power nonlinearity, and normalization.

The linear weights capture the first-order aspects of a given MT neuron’s tuning: its tuning preferences and a coarse estimate of tuning breadth. In fact, both frequency- and velocity-separable

(“linear”) models can capture single grating tuning curve shape well. This is why the two separable models are indistinguishable, on average, when fit to the single grating dataset (figure 5). The model captures the continuum of selectivity for pattern motion in MT (characterized by a unimodal constant frequency plaid direction tuning curve, see figures 8(d) and 11(b)) by simply increasing the direction tuning bandwidth in the linear weights (Simoncelli and Heeger, 1998). Direction tuning bandwidth (calculated from measured tuning curves) is correlated with pattern index, as observed in our data (Pearson’s *r* = 0.27, *P* = 0.0027, n=112) and previous studies (Pearson’s *r* = 0.35, *P <* 0.0002, n=788, Wang and Movshon (2016)). Linear weighting alone in the velocity-separable model is sufficient to capture the unimodality of constant frequency plaid (pattern) direction tuning, but not in the frequency-separable model (lightest red and blue traces, respectively, in figure 11(a)). Further, the frequency-separable model severely underestimates constant velocity plaid responses (figure 11(b)). It is important to note that in order for the two separable models to make realistic and distinguishable predictions, temporal frequency tuning data were also included during fitting; all models capture this tuning well (figure 8). Ultimately, the “linear” model fails by overestimating the tuning bandwidth to frequency plaids (figure 11(a)).

**Figure 11.**
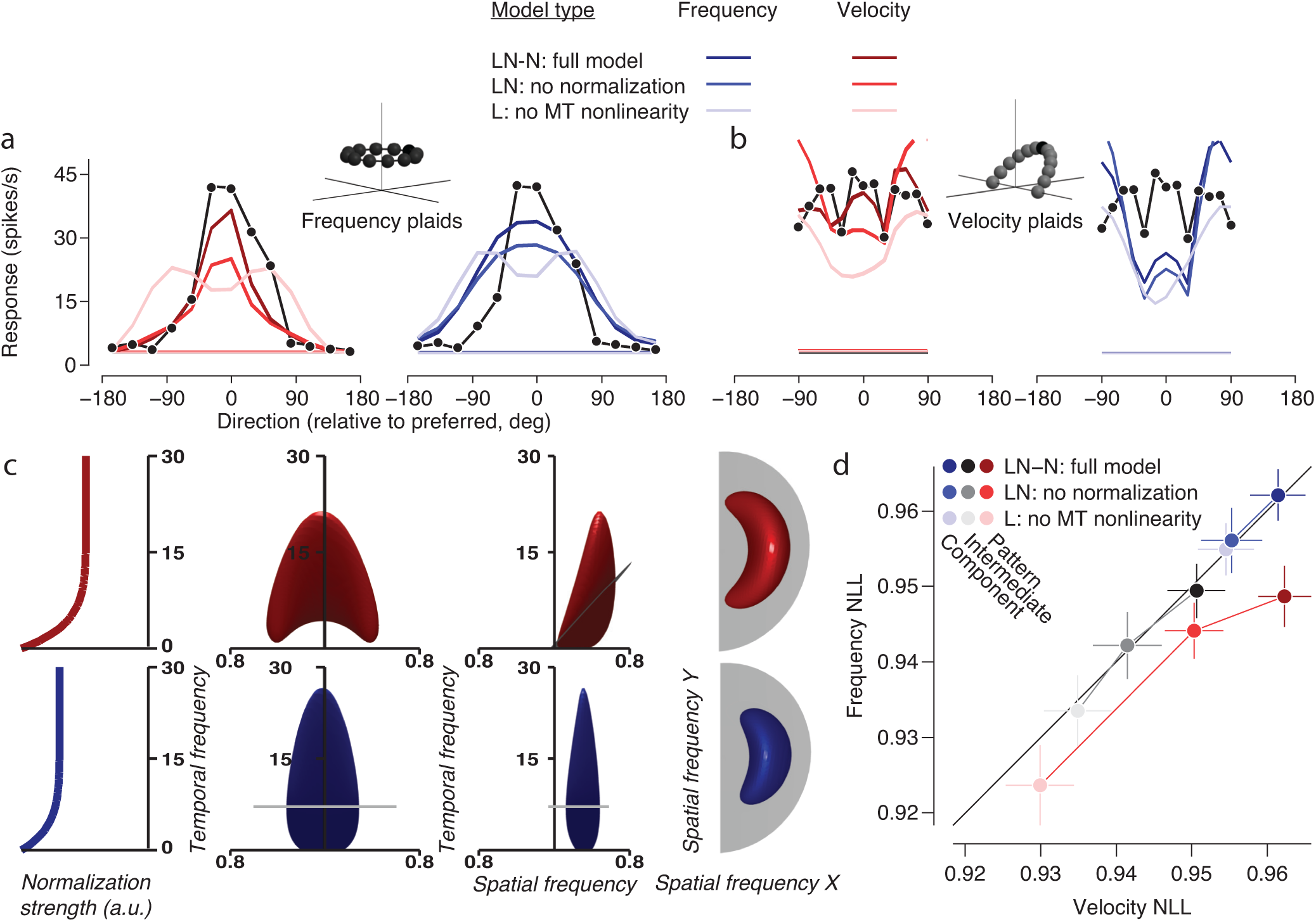
Effects of removing model elements for one example neuron and the population. (a-c) Plots and renderings are from fits to the same neuron shown in figure 8(d). In (a, b, and d), three nested model fits are shown. Lighter shades denote fits with nonlinear elements removed from the full model. LN-N is the full model, which includes linear weighting (L), a point-wise power function nonlinearity (N), and “temporal frequency dependent” normalization (-N). LN has no normalization at all, and L has no nonlinearity at the MT stage. Direction tuning data are shown as black points and lines for constant-frequency plaids (a) and constant-velocity plaids (b). The red and blue shaded curves show the different model fits to those data, with red and blue corresponding to the velocity and frequency models. The darkest traces are for the full model, the lighter ones for the model with normalization removed, and the lightest for the model with no MT nonlinearity. Each (nested) model was optimized separately. (c) The leftmost plots show the strength of normalization as a function of temporal frequency, for the velocity (red, top row) and frequency (blue, second row from top) models. The middle two renderings show the linear weights (at one level set) as a function of spatial and temporal frequency, at two different viewing angles. The temporal frequency scale in these renderings match that of the normalization plots on the left. The renderings on the right are a “birds-eye” view, showing the same weights as function of the two spatial frequency dimensions. (d) Fit quality, expressed as the normalized log likelihood of the velocity and frequency models, is plotted for all component, intermediate, and pattern neurons (in blue, black, and red, respectively), for the three nested models (lighter shades indicate nonlinear model elements removed, see above).

The original Simoncelli and Heeger (1998) model solved this problem by applying an expansive, point-wise nonlinearity in the MT stage. The nonlinearity, when fit to data, only changes tuning to compound stimuli. By adding a point-wise power function, both separable “linear-nonlinear” (LN) model achieve better, sharper constant frequency plaid tuning (medium blue and red traces in figure 11(a)). Frequency-separable predictions to constant velocity plaids (medium blue in figure 11(b)) however, worsen, since tuning for all mixture stimuli are sharpened by the power function. This model parameter is strongly correlated with measured pattern index (figure 10), indicating its role in pattern motion computation.

If the goal was simply to correctly reproduce direction tuning for constant frequency gratings and plaids, a separable model including linear weighting and a power function nonlinearity alone would be sufficient. However, the unexpected, nonlinear property of MT selectivity that we discovered is that direction tuning bandwidth is wider for constant velocity gratings than for constant frequency gratings (figure 6), but by much less than expected, and by much less than is the case for measured constant velocity plaid tuning (figure 8(d)). In order to account for this small change in grating tuning and large change in plaid tuning, the separable models require normalization at the MT stage.

We used a closed-form approximation of MT normalization, rather than simulating an entire population of MT neurons (as was done in the original Simoncelli and Heeger (1998) model, see methods for a details), to make model fitting tractable. Despite being an approximation, it is a functionally interpretable one. Its primary effect is to suppress responses at low temporal frequencies; its tuning relative to the linear weights is plotted in figure 11(c). Suppression for low temporal frequencies has been observed experimentally (Maunsell and Van Essen, 1983). Incorporating normalization improves contrast gain control (darkest blue and red tuning curves are better scaled to the data in figure 11(a,b)). This normalization also sharpens tuning to both single gratings and conjunctions of gratings by concentrating suppression at low temporal frequencies. In the case of pattern neuron responses, these conjunctions are components consistent with a preferred velocity, so velocity-separable model predictions for constant velocity plaids are appropriately widely tuned (figure 11(b)) and frequency-separable model predictions for constant frequency plaids are too widely tuned (figure 11(a)).

For each nested version of the models, namely, the “linear,” “linear-nonlinear,” and the full model, the velocity-separable version performs better on pattern neurons as a group (figure 11(d)). For component and intermediate neurons, the two types of model are indistinguishable regardless of model version.

### Relationship to previous work

Perrone and Thiele (2002) observed broader temporal frequency tuning than predicted by the Simoncelli and Heeger (1998) model. Their Weighted Intersection Mechanism (WIM) model was able to capture joint spatial and temporal frequency tuning in MT. It employed a weighting function on V1 inputs organized along a common speed, and only responded when two types of V1 inputs, “sustained” and “transient,” had equal response levels. With particular choices of parameters, the WIM model can also simulate pattern direction selectivity (Perrone, 2004). The authors stated that a model with velocity-based tuning at the MT stage, such as in Simoncelli and Heeger (1998), would not be capable of producing realistic spatiotemporal tuning. Here we fit just such a velocity-based separable model directly to pattern motion data and to data simultaneously spanning all three dimensions of frequency space. It achieved similar realism in reproducing tuning in both temporal frequency and pattern motion direction, each recorded from a heterogeneous populations of neurons, without the need for the two specific V1 neuron types.

Priebe et al. (2003) investigated joint tuning for spatial and temporal frequency and pattern motion selectivity in MT. Consistent with our findings, they reported stronger evidence for speed tuning with compound stimuli such as plaids or square-wave gratings, as compared to single sinusoidal gratings. Additionally, speed tuning for single sinusoidal gratings and degree of pattern selectivity were independent. They concluded that speed tuning arises in MT only when multiple spatial frequencies are present. Our findings are consistent with these conclusions, and arise in our velocity-separable model simulations as a result of the MT normalization. Our model additionally predicts that pattern tuning will be correlated with speed tuning for square wave gratings and random dots (Kumano and Uka, 2013; McDonald et al., 2014; Xiao and Huang, 2015).

More recent studies (Nishimoto and Gallant, 2011; Inagaki et al., 2016) have explored MT selectivity in all three dimensions of the frequency domain. Nishimoto and Gallant (2011) used “motion-enhanced” natural movies to visualize 3D spectral receptive fields of individual MT neurons for the first time. These weights followed a simulated V1 population, which performed linear filtering, a compressive nonlinearity, and divisive normalization. They reported weights with excitation organized along a partial ring on the plane, with a gap in the ring occurring at low temporal frequencies. Suppression also appeared as partial rings off the preferred velocity plane, much like the opponent suppression reported in Rust et al. (2006). Inagaki et al. (2016) performed linear regression directly on the frequencies of their stimulus, which was comprised of multiple gratings superimposed spatially and partially overlapping in time. They observed broadly tuned receptive fields at mid- and high temporal frequencies in two pattern cells and observed diffuse suppression off the preferred velocity plane. The absence of excitation at low temporal frequencies observed in both studies provides indirect support for our use of suppression there.

Neither Nishimoto and Gallant (2011) nor Inagaki et al. (2016) directly confirmed that their models could produce pattern tuning, making the connection between the receptive field structure they observed and pattern selectivity harder to interpret. The velocity separable model is able to reproduce pattern tuning in both frequency- and velocity-separable coordinates, while making slightly different predictions of receptive field structure. Pattern cells have excitation on a full ring on the preferred velocity plane, with partially overlapping suppression at low temporal frequencies (figure 11(f)). Including normalization at the V1 stage (Rust et al., 2006; Nishimoto and Gallant, 2011) or using a purely subtractive form of suppression in MT (as in all three aforementioned studies) in the separable models was not sufficient to simultaneously account for the broad tuning observed for constant velocity plaids.

### Conclusions drawn from the separable model

Selectivity to moving patterns is a hallmark of MT response. How does this selectivity arise? Orientation selectivity in V1 provides a useful analogy. There, first-order properties of selectivity to simple stimuli, such as simple/complex classification, can be attributed to linearly weighting of LGN afferents (Reid and Alonso, 1995; Goris et al., 2015). Responses to compound stimuli, however, are likely a result of additional (possibly recurrent) computation within V1. Likewise, basic MT direction selectivity along the component/pattern continuum is can be constructed by appropriate summing of V1 inputs (on a constant velocity plane) and shaped on a per-neuron basis by their own point-wise nonlinearities. Further nonlinear tuning behaviors, such as the different tuning bandwidths for constant velocity gratings and plaids, is likely shaped by recurrent computation within MT.

There is some evidence of recurrent computation shaping pattern motion signals in MT. Using drifting fields of bars, Pack and Born (2001) showed that pattern motion tuning emerges later in a pattern neuron’s response—approximately 70ms after its earliest response to stimulus onset— a result later replicated with sinusoidal gratings and plaids (Smith et al., 2005; Solomon et al., 2011). Further experiments could be done to verify this recurrent computation prediction. If feasible, imaging a population of MT neurons and fitting a population-level model could reveal these recurrent computations, as has been done in V1 (Cossell et al., 2015; Antolík et al., 2016; Klindt et al., 2017). Examining dynamics of tuning to compound stimuli, possibly with whole-cell recording techniques, could also provide empirical evidence regarding the nature of suppression in MT.

While the velocity separable model unifies data and theory regarding tuning for pattern direction and velocity, there is much work to be done to further incorporate other aspects of MT selectivity into the model. The velocity separable model includes rudimentary gain control, and we used stimuli which only had two different contrast values. However, accounting for gain control in MT, and its interactions with pattern motion selectivity, motion opponency, and stimulus size (Britten and Heuer, 1999; Heuer and Britten, 2002), will likely require more experiments, and perhaps inclusion of a full normalization pool at the MT stage.

A strict interpretation of the Simoncelli and Heeger (1998) model predicts broad direction tuning to both constant velocity gratings and plaids, yet we only observed broad tuning to the latter (figures 6 and 8(d)). This is consistent with later findings (Priebe et al., 2003; Priebe et al., 2006) suggesting that some speed tuning in MT is inherited from V1, but that full form-independent speed computation occurs within MT, and is evident only when multiple spatial frequencies are present. Our results further suggest that individual MT pattern neurons always signal motion direction, but only signal speed when it is uniquely specified (i.e., when multiple orientations or spatial frequencies are present).

Finally, our findings change our understanding of the degree to which MT corresponds with motion perception. Consider a single drifting contour or grating, viewed through an aperture. The true direction of motion is inherently ambiguous: any drift direction *±* 90 degrees from normal is a valid interpretation.

Perceptually, however, this so-called “aperture problem” is unambiguously solved: the grating is perceived to be drifting in the direction normal to its orientation (Stumpf, 1911; Todorovíc, 1996; Wohlgemuth, 1911; Wallach, 1935; Marr and Ullman, 1981; Adelson and Movshon, 1982). Previously, it was thought that pattern selective neurons in MT, as a population, would signal a single grating’s drift direction ambiguously (Movshon et al., 1985; Simoncelli and Heeger, 1998). Our findings show MT pattern neurons can unambiguously signal such motion, and that such a population can represent the translational motion of a stimulus regardless of whether it contains a mixture of orientations or a single one. The representation of motion in MT may thus be even closer to perception than previously thought.

## Acknowledgements

We thank Romesh Kumbhani, Michael Gorman, Najib Majaj, and other members of the Movshon Laboratory for help with physiological experiments.

---

**Extended data figure 4-1.**
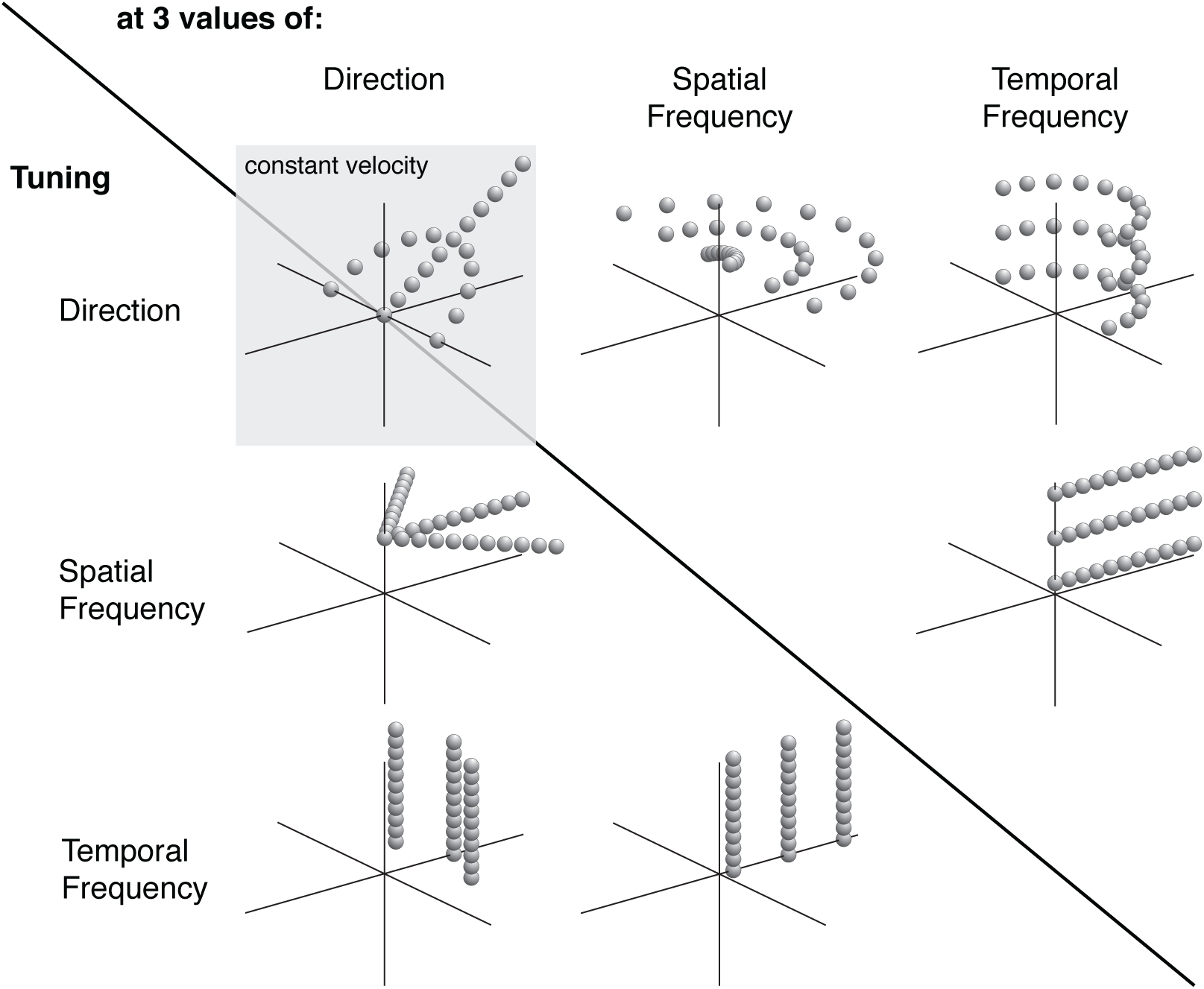
Single grating stimulus set, organized by tuning curves measured relative to preferred values. Top left tuning curves are constant-velocity direction tuning (arc), and constant-velocity spatiotemporal frequency tuning (line). All other tuning curves follow the convention that the type of tuning curve comes from the label on the left, and they are presented at one optimal and two suboptimal values in the dimension derived from the top label. For example, the bottom left tuning curves are temporal frequency tuning curves measured a one optimal direction and two suboptimal ones. Note that optimal tuning curves appear more than once in this figure, but were presented with equal probability during the experiment.

**Extended data figure 4-2.**
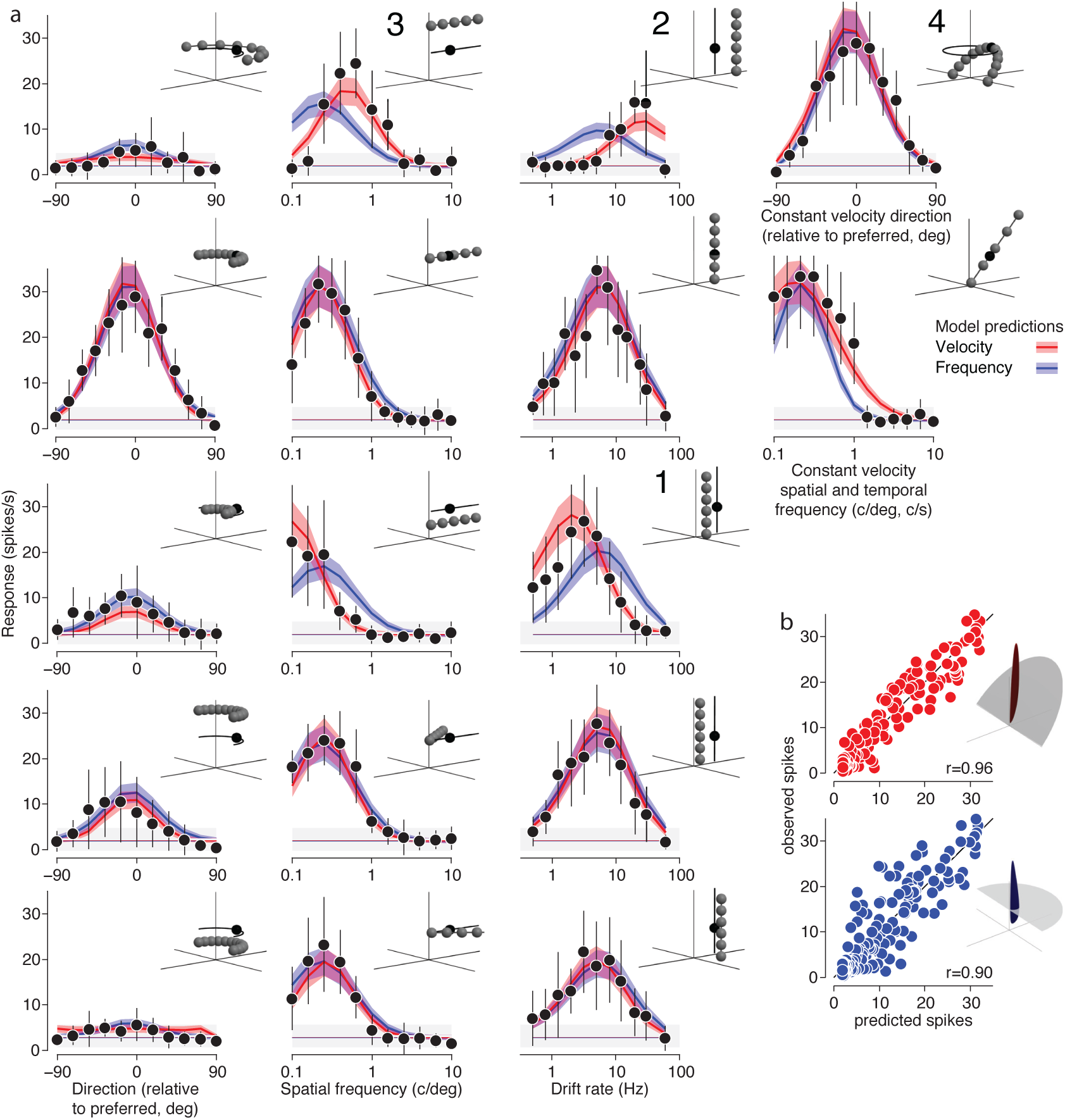
All data and model predictions from the single grating study for the neuron in figure 4(d). (a) All 17 tuning curves in the single grating tuning dataset (see extended data table 4-1 and figure 4-1). Tuning curves marked 1-4 are replicas of the bottom row of tuning curves in figure 4—see its caption for more details. Direction preferences (first column) do not change at different spatial and temporal frequencies, but gain does. Spatial frequency preferences shift at different temporal frequencies (second column, top three rows), but not different directions (second column, bottom three rows). The same is true for temporal frequency preferences (third column). (b) Observed and predicted spikes in response to each of the 225 unique data points in the single grating dataset, for the velocity- and frequency-separable models (red and blue, respectively).

**Extended data table 4-1.**
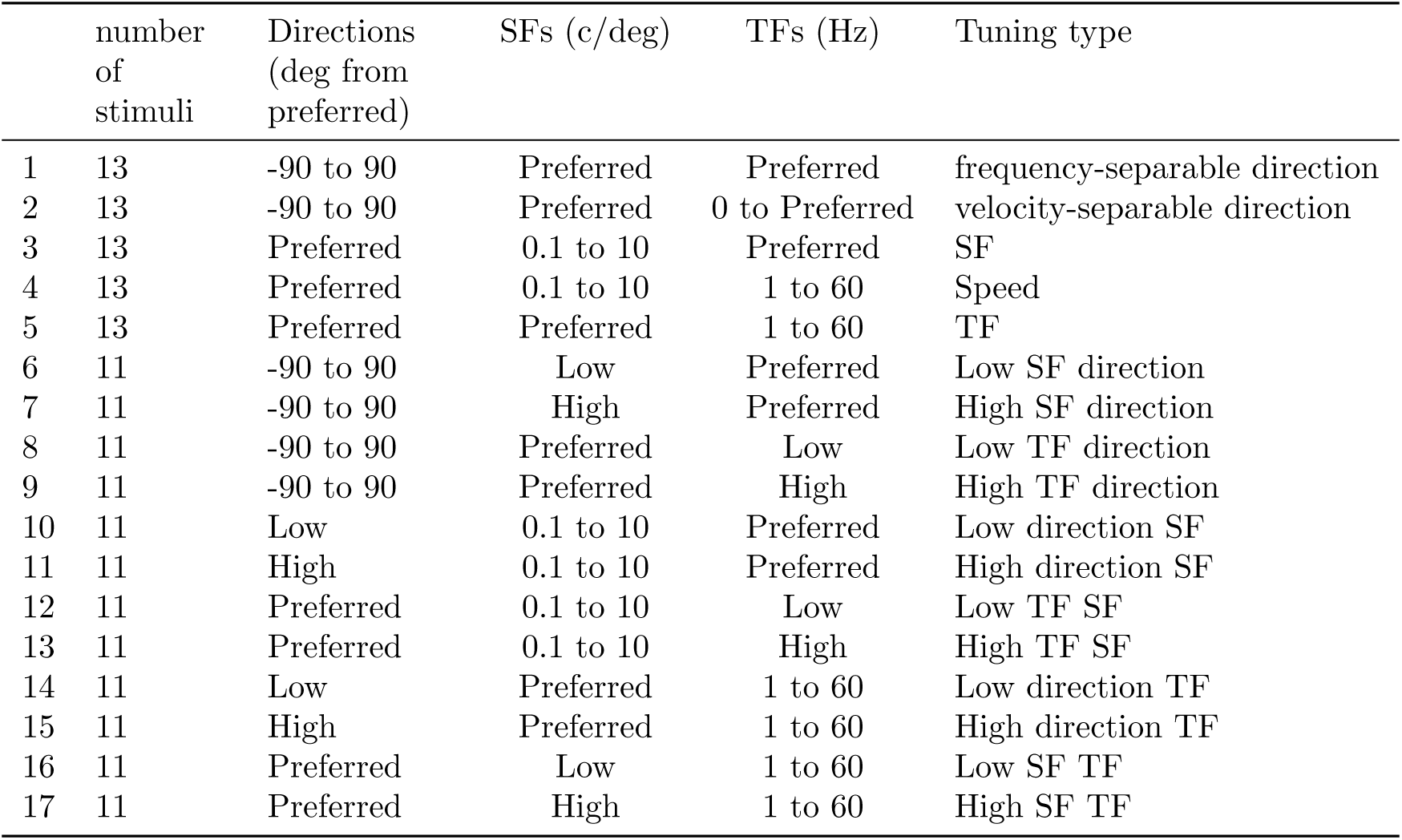
Single grating stimulus set. For the single component study, 17 unique tuning curves were measured, for a total of 225 unique stimulus conditions (figure 4-1 extended data). All featured single gratings presented at 100% contrast. Two direction tuning curves from -90° to 90° relative to the preferred direction, in 15° intervals, were collected along the optimal frequency-separable path (keeping the optimal spatial and temporal frequencies constant) and along the optimal velocity-separable path (keeping the optimal velocity constant). Four direction tuning curves were collected at 18° intervals from -90°to 90° relative to the preferred direction: one at a higher and one at a lower than optimal temporal frequency while fixing the optimal spatial frequency, and two more at a high and a low spatial frequency while fixing the optimal temporal frequency. Two spatial frequency tuning curves, at 13 log-spaced values from 0.1 cycles/degree to 10 cycles/degree, were collected along the optimal frequency- and velocity-separable paths. Four spatial frequency tuning curves, at 11 log-spaced values from 0.1 cycles/degree to 10 cycles/degree, were collected at a high and low temporal frequency while maintaining the optimal direction. Two more were collected at suboptimal directions, while maintaining the optimal temporal frequency. One temporal frequency tuning curve, at 13 log-spaced values from 0.1 cycles/second to 60 cycles/second, was collected at the optimal direction and spatial frequency. Four temporal frequency tuning curves, at 11 log-spaced values from 0.5 cycles/second to 60 cycles/second, were collected at a high and low spatial frequency while maintaining the optimal direction. Two more were collected at suboptimal directions, while maintaining the optimal spatial frequency. The “high” and “low” non-preferred spatiotemporal frequencies used in suboptimal tuning curves were chosen to maximally distinguish the frequency- and velocity-separable models.

## Notes

**Conflicts of interest:** The authors declare no competing financial interests.

